# A single-cell transcriptomics atlas for the parasitic nematode *Heligmosomoides bakeri*: Extrapolating model organism information to non-model systems

**DOI:** 10.1101/2024.02.27.582282

**Authors:** Stephen M. J. Pollo, Hongrui Liu, Aralia Leon Coria, Nicole Rosin, Elodie Labit, Jeff Biernaskie, Constance A. M. Finney, James D. Wasmuth

**Affiliations:** Faculty of Veterinary Medicine, University of Calgary, Calgary, Canada; Host-Parasite Interactions Research Training Network, University of Calgary, Calgary, Canada; Department of Biological Sciences, Faculty of Science, University of Calgary, Calgary, Canada

## Abstract

Single-cell atlases aim to collect the gene expression information for every cell type in an organism but can be challenging to perform in non-model organisms. To try to circumvent the problem of having no verified cell type markers in the parasitic nematode *Heligmosomoides bakeri* to use for an atlas, we attempted to use orthologs of verified markers from the closely related model organism *Caenorhabditis elegans*. This resulted in a useful comparison between the two worms for each of the cell types recovered in preliminary *H. bakeri* single-cell RNA-sequencing. For *H. bakeri* males and females, robustly recovered cell types include the gametes, embryos, and male intestine, while hypodermis, neurons, muscles, and pharyngeal cells were under-represented cell types. The two worms appear to have a similar hypodermis, cuticle, eggshell, and spermatogenesis process. On the other hand, putative cell identities and cell cycle scores suggest the intestine and muscle cells in *H. bakeri* may still be cycling and dividing, unlike in *C. elegans*. Additionally, embryogenesis and early development appear to be quite different between the two worms, with only eight out of 94 confirmed paternal contributions to the embryo in *C. elegans* (with an ortholog) predicted to also be paternal contributions in *H. bakeri*. Overall, this new dataset allowed me to move beyond the presence or absence of orthologs to include their tissue specificity and expression level similarities and differences when comparing these two worms to better identify biological processes and traits in a parasitic nematode that are modelled well by *C. elegans*.

## Introduction

Bulk RNA sequencing (RNA-seq) provides valuable gene expression information for populations of cells. However, because the cells are lysed together, the expression information obtained represents the average expression of each gene across the population of cells. Information on the variance in expression of each gene between individual cells is lost. This is particularly relevant in samples with multiple cell types, like tissue samples or whole organisms, because different cell types can have drastically different gene expression from each other (Chen, Teichmann & Meyer, 2018). Single-cell RNA sequencing (scRNA-seq) addresses this loss by dissociating the sample into a single-cell suspension and profiling the cells at single-cell resolution. The resulting data can be used to cluster the cells based on their gene expression profile in order to try to identify the cell types captured (Luecken & Theis, 2019). Collecting the gene expression profiles in this way for all of the cell types in an organism is referred to as a single-cell atlas (Chen, Teichmann & Meyer, 2018). They deepen our understanding of an organism, both through the discovery of previously unknown cell types or cell states, as well as by unravelling the different activities of the various cell types that make up the organism. This is exemplified in the atlas for the planarian *Schmidtea mediterranea*, through which Fincher and colleagues uncovered a novel cell type that was distributed throughout the body with long processes into the parenchymal space (Fincher et al., 2018). They also found an important function of the muscles includes expressing genes that convey positional information throughout the worm.

Of the multiple methods available for generating scRNA-seq data, the 10X Genomics Chromium system offers convenience and an acceptably high capacity for cells profiled per run. Using this system, the typical workflow for generating scRNA-seq data begins with collecting cultured cells or by dissociating tissues into a single-cell suspension. The cell suspensions are then loaded into the Chromium, which aims to capture, within a single droplet, one cell and a gel bead that contains the oligonucleotides needed to profile the RNA within the cell (Zheng et al., 2017). The nature of the oligonucleotides within the bead allows for barcoding of RNA that is released during lysis of the cell within the droplet, as well as barcoding of the exact mRNA that is captured by the poly-dT component of each oligo (Zheng et al., 2017). The captured RNA is reverse transcribed and prepared into a library suitable for sequencing on an Illumina machine. However, the simultaneous barcoding and capturing of the mRNAs by their poly-A tail, along with the length of sequencing possible on the Illumina platform that is used at the end of the process, means that the transcriptome of the cell is profiled at the 3’ end of the transcripts only. In contrast to bulk RNA-seq profiling of the full transcripts, this means there is little to no information on alternative isoforms and overall intron-exon structure. Moreover, approximately 30% of the transcripts are captured per cell (Zheng et al., 2017), meaning that the most abundant cellular transcripts are more likely to be represented in the final library, whereas less abundant transcripts may be missed entirely. This limitation is partially compensated for by profiling many cells of the same type to collectively get a more comprehensive view of the expression patterns of that cell type (Zheng et al., 2017).

After sequencing, demultiplexing the reads using the barcodes allows for identification of which reads came from which cell and for removal of any PCR artifacts (Zheng et al., 2017). The reads are mapped to a reference genome and annotation to generate a matrix of read counts per gene in each profiled barcode. Downstream analyses vary from this point, but for an atlas, the barcodes are clustered based on their expression profiles in order to yield clusters of the different cell types recovered (Luecken & Theis, 2019). The cell identities of the clusters are determined based on the expression of verified cell type markers and/or by hybridization experiments targeting genes found to be up-regulated in clusters of interest (Luecken & Theis, 2019).

Doing this process in model organisms that have high quality genome assemblies and manually curated annotations—like the human and mouse data used to verify the Chromium system—works very well (Zheng et al., 2017). In non-model organisms, however, lower quality genome assemblies may interfere with the ability to map the reads, causing real data to be discarded. Additionally, low quality annotations (missing genes, incorrect intron-exon predictions, incorrect stop coordinates) can cause properly mapped reads to not be included in downstream analyses because they cannot be confidently assigned to a gene in the matrix. While the software used to process 10X Genomics data, CellRanger, has some flexibility to try to compensate for incorrect gene models, missing genes will always cause data loss. Additionally poor-quality functional annotation hinders downstream interpretation of results, and a lack of verified cell type markers complicates identification of the recovered cell types.

One justification for studying model organisms and model systems is that the insights gained can be extrapolated to other systems of interest that are harder to study. Indeed, many processes have been discovered in *Caenorhabditis elegans* that have been found to also occur in other organisms, including RNA interference and developmental apoptosis pathway(s) (Ellis & Horvitz, 1986; Fire et al., 1998). As such, *C. elegans* has been, and continues to be, a useful model animal and should surely be an even better model for nematodes. However, for parasitic nematodes, that have human health, veterinary, or economic relevance, a main concern for using *C. elegans* as a model is that it is not a parasite (Blaxter, 1998). Consequently, traits associated with parasitism (ex. mode of feeding, exposure to host(s) environment and immune system, migrations within/between hosts) may not be modelled well in *C. elegans* (Gilleard, 2004). Moreover, with increasing phylogenetic distance, fewer traits and molecular functions would be expected to be shared (Geary & Thompson, 2001; Gilabert et al., 2016). Therefore, comparisons between *C. elegans* and parasitic nematodes are needed to uncover the similarities and differences between the two, not only in gene content, but also in gene expression and molecular function, to better understand how to best extrapolate knowledge from *C. elegans* to parasitic nematode systems.

The murine intestinal roundworm *H. bakeri* is a non-model organism with a publicly available genome assembly and automated annotation (Chow et al., 2019). Though there are no verified cell type markers for this organism, it is closely related (383 MYA divergence time) to the model organism *C. elegans* (Smythe, Holovachov & Kocot, 2019), for which there are many verified and characterized cell markers. Moreover, orthologs have been pre-computed between the two worms, using a standard pipeline based on gene trees (Vilella et al., 2009), and are available from the resource WormBase ParaSite (Howe et al., 2017). We opted to use the pre-computed orthologs of known *C. elegans* cell markers to attempt to circumvent the problem of having no markers to use to identify cell types in *H. bakeri* scRNA-seq clusters. Doing so presents an inherent comparison between the two worms at the level of gene expression in each of the cell types recovered, especially since multiple scRNA-seq atlases exist for *C. elegans*, including for adults (Ghaddar et al., 2022), the L2 larval stage (Cao et al., 2017), and embryos (Packer et al., 2019).

Here, we have analyzed data from preliminary attempts to generate a single-cell atlas for *H. bakeri*. While the number of cells recovered is insufficient for a complete atlas, and questions remain surrounding sample processing, the use of orthologs of genes in *C. elegans* in the analysis uncovered similarities and differences between the two worms. In particular, moving beyond the presence or absence of orthologs to including their tissue specificity and expression level similarities and differences helps to better identify biological processes and traits that are or are not shared between the two worms. This may serve as a useful case study for the applicability of *C. elegans* biology to other parasitic nematodes, particularly those classified into Clade V.

Note: A version of this preprint was first made available as a chapter in SMJP’s PhD thesis (UCalgary Vault https://hdl.handle.net/1880/117625).

## Materials and Methods

### Mice and parasites

Male C57Bl/6 mice aged 8-10 weeks (bred and maintained at the animal care facility, Department of Biological Sciences, University of Calgary) were used. All animal experiments were approved by the University of Calgary’s Life and Environmental Sciences Animal Care Committee (protocol AC17-0083). All protocols for animal use and euthanasia were in accordance with the Canadian Council for Animal Care (Canada). Infected mice were orally gavaged with 400 third stage *H. bakeri* larvae and euthanized at 10 days post initial infection. Worms were removed from the intestinal tract and placed in Dulbecco’s modified eagle’s medium – high glucose (Sigma cat. D5796) where they were separated by sex.

### Worm dissociation for samples CF4 and CF5

Collected worms were washed in 10% gentamicin for 20 minutes to eliminate bacterial contamination. Worms were placed in a digestion solution consisting of DMEM with DNAse I (0.05%) and Liberase (2%) and incubated in a shaker at 37°C for 30 minutes. The content was then passed through a 40 µm filter and cells were spun down at 1,500 rpm for 5 minutes twice. Cells were then stained with FVS780 (BD, #565388) for 15 minutes at room temperature and rinsed with HBSS with 2% BSA. Cells were then stained with Vybrant DyeCycle 1:500 (ThermoFisher, #V35004) for 1 hour at room temperature.

### 10X Genomics library preparation and sequencing

Approximately 12,000 single cells from each sample were loaded for partitioning using 10X Genomics NextGEM Gel Bead emulsions (v3.1). Each sample was processed according to the manufacturer’s recommended protocol (PCR amplification steps were run at 12X, and 14X respectively). Final cDNA library size determination and QC was performed using TapeStation D1000 assay. Sequencing was performed using Illumina NovaSeq S2 and SP 100 cycle dual lane flow cells over multiple rounds at the UCalgary Centre for Health Genomics and Informatics (CHGI).

### Data availability

All sequence data was deposited in the SRA under the accession number PRJNA1009113.

### Preparation of genome annotation references and mapping and quantification of scRNA-seq data

scRNA-seq data were processed using the 10X Genomics analysis pipeline CellRanger v 7.0.1. To construct the CellRanger reference from the Wormbase ParaSite genome and annotation files for *H. bakeri* (PRJEB15396), the annotation gff3 file had to be modified to replace ‘ID’ and ‘Parent’ tags and change other formatting in field 9. The CellRanger mkref command was then able to construct the references for further analysis. Counts for each barcode in each library used for preliminary analyses were generated using the CellRanger count command. The mapping files generated in the outputs were then merged and sorted into a single bam file for all 10X Genomics data generated in this study. This file was used to extend the annotations to include 3’ UTRs using the program peaks2utr v 0.5 (Haese-Hill, Crouch & Otto, 2023). A modified annotation file to create unique CDS lines for every transcript was required for peaks2utr to run. The resulting annotation file was modified as above to prepare it for CellRanger mkref. CellRanger count was then run on all libraries with the new reference to map and count the reads for each barcode. The raw feature barcode matrices were then downloaded into R for further analysis.

### Construction of male and female single-cell atlases

#### Preliminary analyses

Preliminary analyses were conducted with the CellRanger pipeline. The libraries analyzed with CellRanger were merged into a single analysis with the CellRanger aggr command. The results of the full pipeline, including cell clustering and marker gene analysis were visualized and explored in the 10X Genomics Loupe browser v 6.2.0.

#### Quality control and filtering of single-cell data

Raw feature barcode matrices from CellRanger were processed with the R package Seurat v 4.0.2 (Hao et al., 2021), following the vignettes for Guided Clustering, Cell-Cycle Scoring and Regression, and Introduction to SCTransform, v2 regularization (https://satijalab.org/seurat/index.html). Briefly, for the males and females separately (See Results and Discussion for preliminary analysis on a single merged atlas), Seurat objects were created from the raw feature barcode matrices and merged into a single object. The object was filtered according to the number of features each cell had to remove empty barcodes and potential doublets. Counts of mitochondrial features could not be obtained because the reference genome available at this time does not contain mitochondrial sequence to map to, and therefore no annotated mitochondrial genes.

#### Data normalization and clustering

Each Seurat object was initially processed following the vignette for Guided Clustering, using log normalization, separate selection of 5000 variable features, linear scaling, and PCA dimensional reduction. This was done to enable calculation of cell cycle scores with the Seurat CellCycleScoring method, which would not work on the SCT slot values of objects processed with the newer SCTransform method. Once cell cycle scores were calculated, the SCTransform method with v2 regularization was used to re-normalize the original count data while regressing out the cell cycle scores. PCA was then used for dimensionality reduction, followed by uniform manifold approximation and projection (UMAP) for visualization. Clusters were determined using the FindNeighbors method with 30 dimensions and the FindClusters method with the default resolution of 0.8, both from Seurat.

### Marker genes, *C. elegans* orthologs, and putative cluster annotation

Marker genes were calculated for every cluster in both atlases using the Seurat method FindAllMarkers. Features that were up-regulated in the cluster by at least a log-fold difference of 0.25 relative to the rest of the atlas and were detected in a minimum of 25% of cells in either the cluster or the rest of the atlas were retained as markers. Since there are no verified cell or tissue markers in *H. bakeri*, we aimed to use orthologs of known markers in *C. elegans*. All orthologs between *H. bakeri* and *C. elegans* were retrieved from WormBase ParaSite (8298 unique *H. bakeri* genes) (Howe et al., 2017). Individual markers or combinations of markers in *C. elegans* were obtained from literature, including wormbook, wormatlas, and individual papers found during literature searches. When multiple markers for a tissue type were collected, a module score was calculated for every cell in each atlas using the AddModuleScore method in Seurat. Module scores and key marker genes were used to putatively annotate the clusters in the atlases.

#### Sources of marker gene modules

The first study sequenced from mixed-stage *C. elegans* worms RNA that was bound to a polyA-binding protein that was expressed under the control of different tissue-specific promoters, including *ges-1* for intestinal expression, *myo-2* for pharyngeal muscle expression, and *myo-3* for body wall muscle expression (Blazie et al., 2015). By comparing these datasets to each other, they defined transcripts that were uniquely expressed in each tissue, relative to the others, as genes with a fragments per kilobase per million mapped reads (FPKM) ≥ 1 in that tissue but undetected (or FPKM < 1) in the other tissues. This resulted in 4091 unique intestinal genes, 312 unique pharyngeal muscle genes, and 329 unique body wall muscle genes. The modules of *H. bakeri* orthologs of these genes are referred to here as Blazie_int_uniq, Blazie_pharynx_uniq, and Blazie_bodymuscle_uniq, respectively.

The second study performed RNA-seq on synchronized *C. elegans* embryos every 30 minutes starting at the 4 cell stage (Boeck et al., 2016). Through their online portal (GExplore - http://genome.sfu.ca/gexplore) we were able to perform comparisons between various timepoints to define gene sets of interest, as well as retrieve some of their pre-defined sets like the maternally enriched set of genes (matenr). Importantly, their later timepoints (122 minutes or more) represent embryonic stages that are post-egg-laying but pre-hatching. Modules resulting from comparisons of this data set are named Boeck_gex_[set description], where pre-defined sets retain their name (matenr, afterlayingenr) and comparisons we performed are described (ex. Ya_g10_83_g10_277_g10_44_g10_161 is genes that are up-regulated in young adult worms by more than 10 fold compared to the 83 min sample and the 277 min sample and the 44 min sample and the 161 min sample).

The third study sequenced 3’ ends of transcripts within intestinal nuclei of mixed-stage *C. elegans* worms that were obtained from fluorescence-activated nuclei sorting (Haenni et al., 2012). They compared a sample of sorted intestinal nuclei to an unsorted sample to identify 2456 genes with higher expression in the intestinal nuclei. The module of *H. bakeri* orthologs of these genes is referred to here as Haenni_intestine.

The fourth study performed bulk RNA-seq on sorted cell populations from adult *C. elegans* worms (Kaletsky et al., 2018). They sequenced hypodermal cells using a *pY37A1B.5::gfp* reporter strain, intestinal cells using a *Pges-1::gfp* reporter strain, neurons using a *Punc-119::gfp* reporter strain, and body muscle cells using a *Pmyo-3::mCherry* reporter strain. Comparing amongst their samples allowed them to define genes that were enriched in (highly expressed and significantly differentially expressed relative to the average expression of all the other tissues) or unique to (highly expressed and significantly differentially expressed relative to each of the other tissues) each tissue. Each of the modules of *H. bakeri* orthologs of these genes are referred to here as Kaletsky_adult_[tissue]_[enriched OR unique].

The fifth study performed bulk RNA-seq on sorted cells from synchronized *C. elegans* embryos of several fluorescent reporter strains including muscle (and coelomocytes; *hlh-1p::mCherry*), intestine (*end-1p:: mCherry*), neurons (*cnd-1p::mCherry*), pharynx (*pha-4::GFP*), and hypodermis (*nhr-25::GFP*) (Warner et al., 2019). They sampled every 90 minutes starting at egg laying for five timepoints. By clustering genes according to their expression patterns throughout the tissues and timepoints, they were able to define genes important for different embryonic tissues, as well as genes that were broadly expressed and decreasing from the point of egg laying. Each of the modules of *H. bakeri* orthologs of these genes are referred to here as Warner_embTS5_[warner cluster].

### Average cluster profiles, GO enrichment, and differential gene expression analysis

Average expression profiles were calculated from the final atlases for each cluster using the AverageExpression method in Seurat. Gene ontology (GO) enrichment was performed using the R package gprofiler2 v0.2.1 (Kolberg et al., 2020). Differential gene expression analysis was conducted in Seurat using the FindMarkers method. For male intestinal comparisons, clusters 1, 3, 7, 8, and 10 were each compared to the remaining clusters in the atlas. The intersections of these comparisons were computed in order to explore features that were statistically significantly (padj < 0.05) up- or down-regulated in all intestinal clusters relative to non-intestinal tissues. To compare between the intestinal clusters, all pairwise comparisons were performed between clusters 1, 3, 7, 8, and 10.

### Cross-species analysis with LIGER

A cross-species analysis between scRNA-seq data from adult *C. elegans* hermaphrodites (Ghaddar et al., 2022) and the *H. bakeri* data set generated in this work was performed with the R package LIGER v 1.1.0 (Welch et al., 2019). *H. bakeri* datasets were read into R using the Read10X method from the Seurat package v 4.0.2 (Hao et al., 2021). Barcodes were given unique names so the sparse matrices could be merged using the RowMergeSparseMatrices method from Seurat. Each Seurat object was then filtered to remove empty barcodes and potential doublets. The three *C. elegans* datasets as sparse matrix .rds objects were read into R and merged using the RowMergeSparseMatrices method from Seurat. This object was filtered to remove empty barcodes and potential doublets and merged with the *H. bakeri* Seurat object. The resulting Seurat object was then converted to a LIGER object with the seuratToLiger method in LIGER. The analysis was repeated with different combinations of included *H. bakeri* data: 1) all the male *H. bakeri* libraries were included, 2) all the female *H. bakeri* libraries were included, or 3) all the *H. bakeri* libraries were included.

A requirement of the LIGER analysis that went unmentioned in the documentation is that shared features (genes) between the species being compared must have the same name in the raw dataset. To accomplish this, the orthologs between *H. bakeri* and *C. elegans* identified above were searched against the total list of features in the *H. bakeri* data within the LIGER object to replace the names of the orthologous genes in *H. bakeri* with the locus tag being used for the *C. elegans* gene. The remaining analysis was then conducted following the LIGER vignette for “Cross species Analysis with UINMF” (http://htmlpreview.github.io/?https://github.com/welch-lab/liger/blob/master/vignettes/cross_species_vig.html), where thresholds were set to 0.3 and the *H. bakeri* sets were allowed to have unshared features while selecting genes. For the optimizeALS step lambda was set to 5, k was set to 30, and the threshold was 1e-10. Finally, the *C. elegans* datasets were set as the reference during the quantile_norm step.

To relate the Seurat results above to the LIGER results to the results from (Ghaddar et al., 2022), a table was constructed for all barcodes in the *C. elegans* and *H. bakeri* datasets. The cluster assigned to each cell in each analysis, along with the cluster annotation from (Ghaddar et al., 2022) for the *C. elegans* cells was included. The proportion of *C. elegans* cells of each assigned cell type or cell group was weighted by the proportion of the LIGER cluster in each Seurat cluster and summed to get putative cell types of Seurat clusters based on the combined *H. bakeri*/*C. elegans* analysis.

## Results and Discussion

### Libraries and cells

Six 10X libraries were prepared and sequenced by collaborators (Table 1). Samples were sequenced to > 40,000 reads per cell (> 50% sequencing saturation). Across the samples 800 – 5800 cells were captured (after filtering), with the exact number of cells recovered in each library after filtering in Table S5. This depth of sequencing resulted in 1200 – 2000 median genes per cell.

**Table 1.** Sample metadata.

| Sample Name | Worm sex | Batch | Dissociation enzyme |
| --- | --- | --- | --- |
| CF4 | M | 1 | Liberase |
| CF5 | F | 1 | Liberase |
| ML | M | 2 | Liberase |
| MP | M | 2 | Pronase |
| FL | F | 2 | Liberase |
| FP | F | 2 | Pronase |

### 3’ profiling with 10X Genomics Chromium defines many 3’ UTRs

The existing annotation for *H. bakeri* from WormBase ParaSite has annotated 3’ UTRs for only 47% of the transcripts. Using the program peaks2utr to extend the annotated regions to include 3’ UTRs based on the mappings of the 10X Genomics data enabled 14449 3’ UTRs (57% of all transcripts) to be annotated, including extending existing 3’ UTRs (Table 2). Considering that the 10X data is specifically profiling the 3’ ends of the transcripts, while the transcript ends are underrepresented in bulk RNA-seq data (Wang, Gerstein & Snyder, 2009), these new predictions are preferable. These newly predicted 3’ UTRs in *H. bakeri* will undoubtedly be helpful for further investigations into gene expression in this organism, since 3’ UTRs are known to contain elements that regulate gene expression post-transcriptionally (Bartel, 2009). Moreover, tissue-specific UTRs in *C. elegans* have been shown to contain important microRNA targets (Blazie et al., 2015), which may serve as important comparisons to the closely related *H. bakeri*.

**Table 2.**
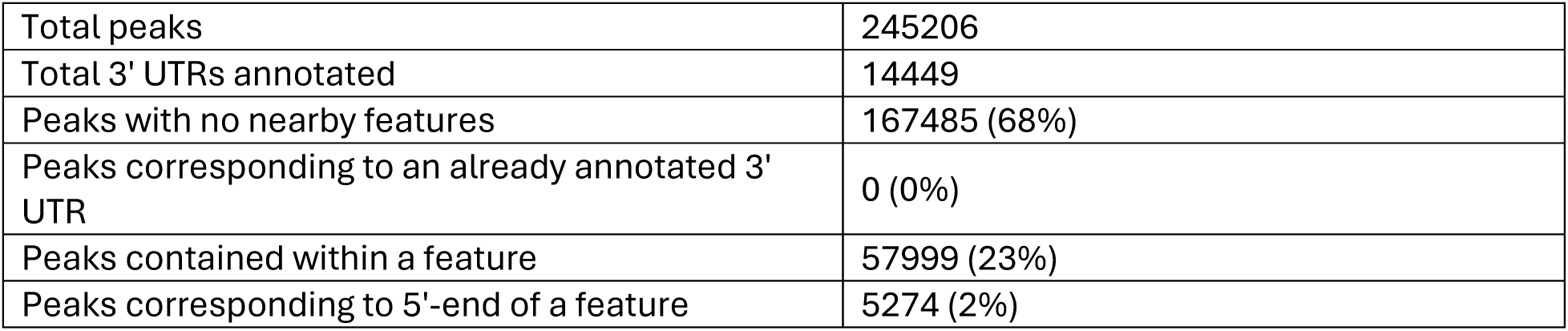
Results from peaks2utr.

|  |  |
| --- | --- |
| Total peaks | 245206 |
| Total 3' UTRs annotated | 14449 |
| Peaks with no nearby features | 167485 (68%) |
| Peaks corresponding to an already annotated 3' UTR | 0 (0%) |
| Peaks contained within a feature | 57999 (23%) |
| Peaks corresponding to 5'-end of a feature | 5274 (2%) |

### Preliminary analyses of a combined male and female single-cell atlas show unexpected major differences between datasets

A preliminary analysis using the 10X Genomics CellRanger pipeline to pool all the H. bakeri libraries into one single-cell atlas was conducted. This enabled me to look for batch effects between the libraries and samples and to determine if pooling all the cells provided additional information to better resolve different tissues into different clusters. UMAPs of the resulting atlas are shown in Figure 1. While no obvious batch effects are apparent here from the use of liberase vs pronase during cell dissociation, or between libraries of the same worm sex, a pronounced batch effect is seen between the male and female worm samples (Figure1D). In our previous analysis of bulk RNA-seq of whole worms (Pollo et al., 2023), the male and female worms at 10 days post-infection (the same time point as used here) were found to statistically significantly differently express 70% of their transcripts. Part of that is due to the different gametes each sex produces and differences in gene expression related to reproduction. Nonetheless, near complete segregation of the male and female cells from each other was unexpected, particularly for non-reproductive tissues that are found in both sexes (ex. Intestine, body muscle, hypodermis, etc.). It is unclear from this analysis whether the differences between the male and female samples reflect: 1) significantly different transcription profiles in all tissues of each sex, to the point that cells of the same tissue type do not cluster together in unsupervised clustering approaches, or 2) a significant difference in the cell types recovered in the male and female samples, to the point that little to no overlap was recovered between the two. To investigate these possibilities, and to have the clusters, and resulting transcription profiles, reflect cell state differences rather than sample differences, we performed the rest of the analyses on separate male and female atlases that were constructed and analyzed in parallel.

**Figure 1.**
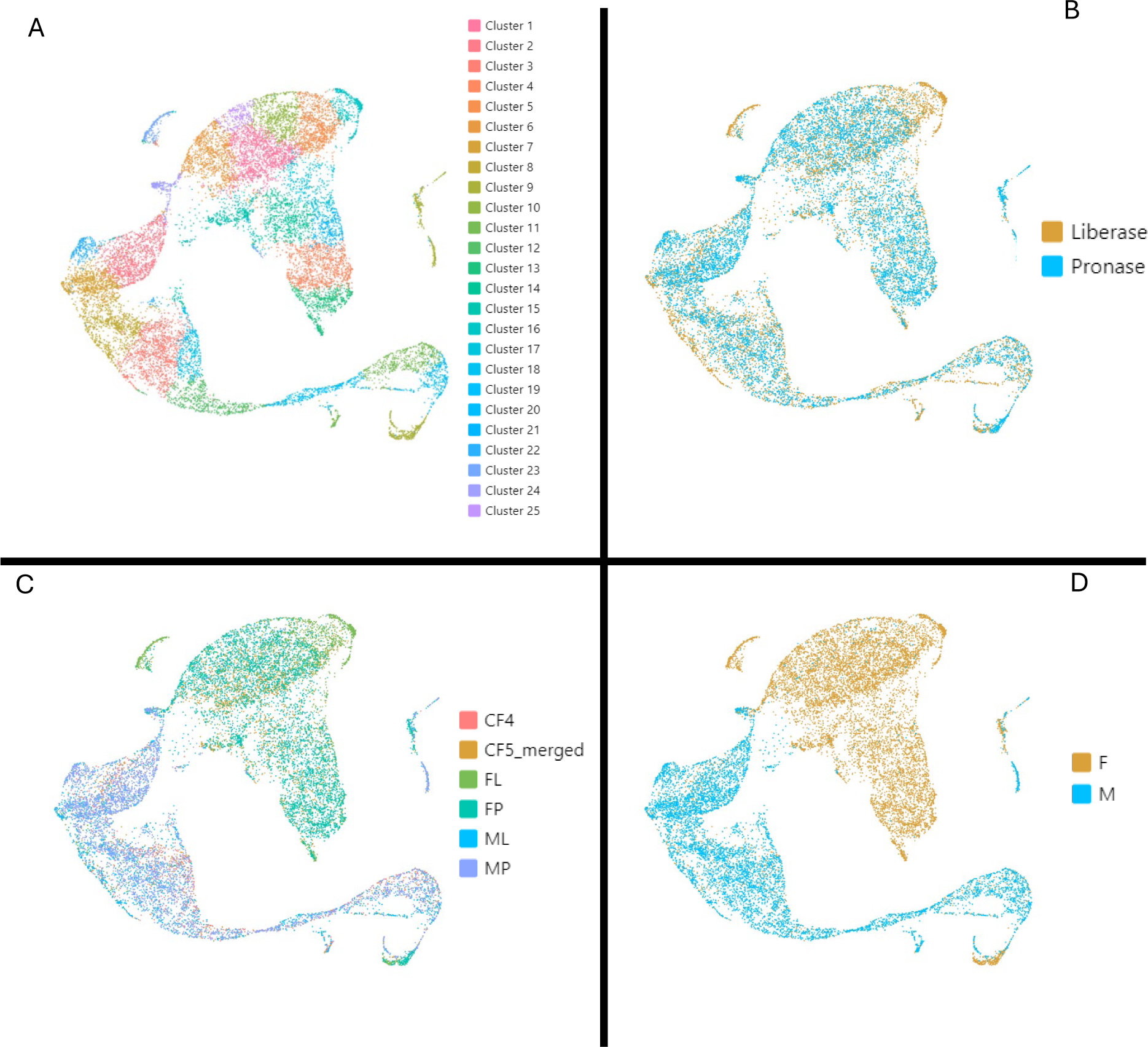
UMAP representation of the preliminary single-cell atlas of combined male and female samples. Each cell of the atlas is coloured according to A) the clusters assigned by the CellRanger pipeline, B) the enzyme cocktail used during cell dissociation, C) the sample library, or D) the sex of the worms in the sample.

### Separate male and female single-cell atlases and putative cluster annotations

#### Final male and female atlases show more even distribution of cell cycle state and UMI counts across the atlas

Preliminary atlases had clusters driven by cell cycle state, and/or affected by UMI count (Figure S1). By regressing out cell cycle scores during normalization and using the updated Seurat v2 regularization method of the new SCTransform method, the final atlases have a more even distribution throughout the clusters of cells in different phases of the cell cycle and cells with different levels of UMI counts (Figure 2). Additionally, visualizing the expression levels of features considered to be cluster markers often highlights individual clusters rather than showing general high expression throughout the atlas (See for example Figure S2).

**Figure 2.**
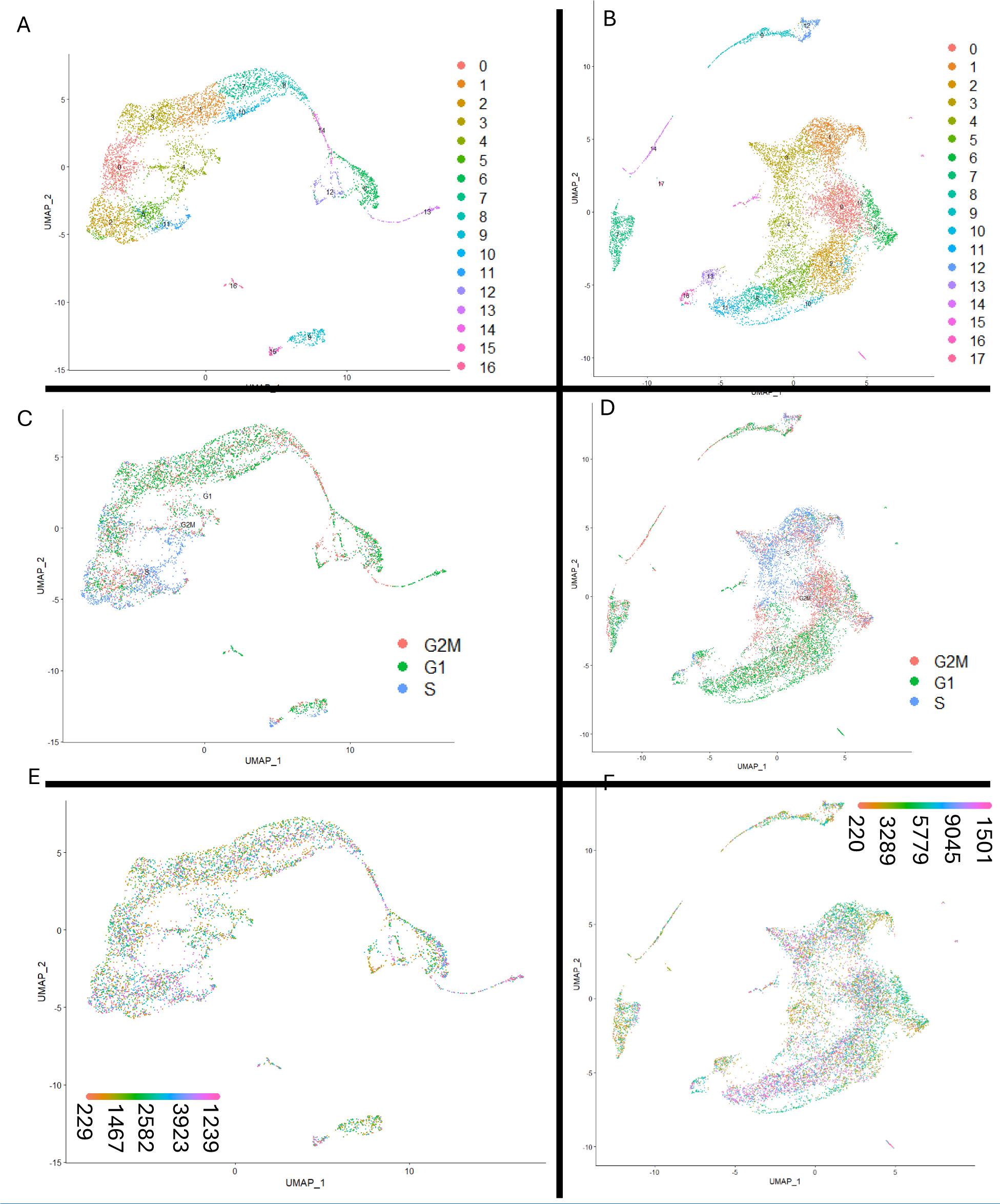
UMAPs of the final single-cell atlases. A, C, and E are the male atlas. B, D, and F are the female atlas. A and B show cells coloured by assigned cluster, C and D show cells coloured by assigned cell cycle phase, and E and F show cells coloured by UMI count.

The clusters in the male and female atlases (See Table S5 for stats on clusters and cells) were putatively annotated on the basis of cell types implicated by marker genes for the cluster and/or the expression of orthologs of genes in *C. elegans* that are known to identify certain cell types. Similar to the cell cycle scoring method in Seurat, when large collections of genes for a certain cell type were used, the genes were treated as a ‘module’ for which a module score was calculated, where higher scores reflect an enrichment in expression of the module genes in a particular cell relative to a set of random genes.

#### Annotating clusters: putative sperm

As reviewed in (L’Hernault, 2006) and shown in Figure 3, in *C. elegans* males and L4 hermaphrodites, spermatogenesis begins with a syncytium of germ cells connected to a shared cytoplasmic core. Individual primary spermatocytes bud off and proceed through meiosis I to become secondary spermatocytes. As the secondary spermatocytes proceed through meiosis II to become spermatids, they divide in such a way that all ribosomes get left behind in a shared residual body, while organelles like the nucleus, mitochondria, and fibrous body-membranous organelles (FB-MOs) go into the spermatids. FB-MOs contain major sperm protein, which is integral for the pseudopod-based motility of the final sperm. The spermatids lack tubulin, actin, ribosomes, and most voltage-gated ion channels, and are what get released from the male during mating. Final development of male spermatids into mature spermatozoa occurs within the hermaphrodite uterus prior to fertilization.

**Figure 3.**
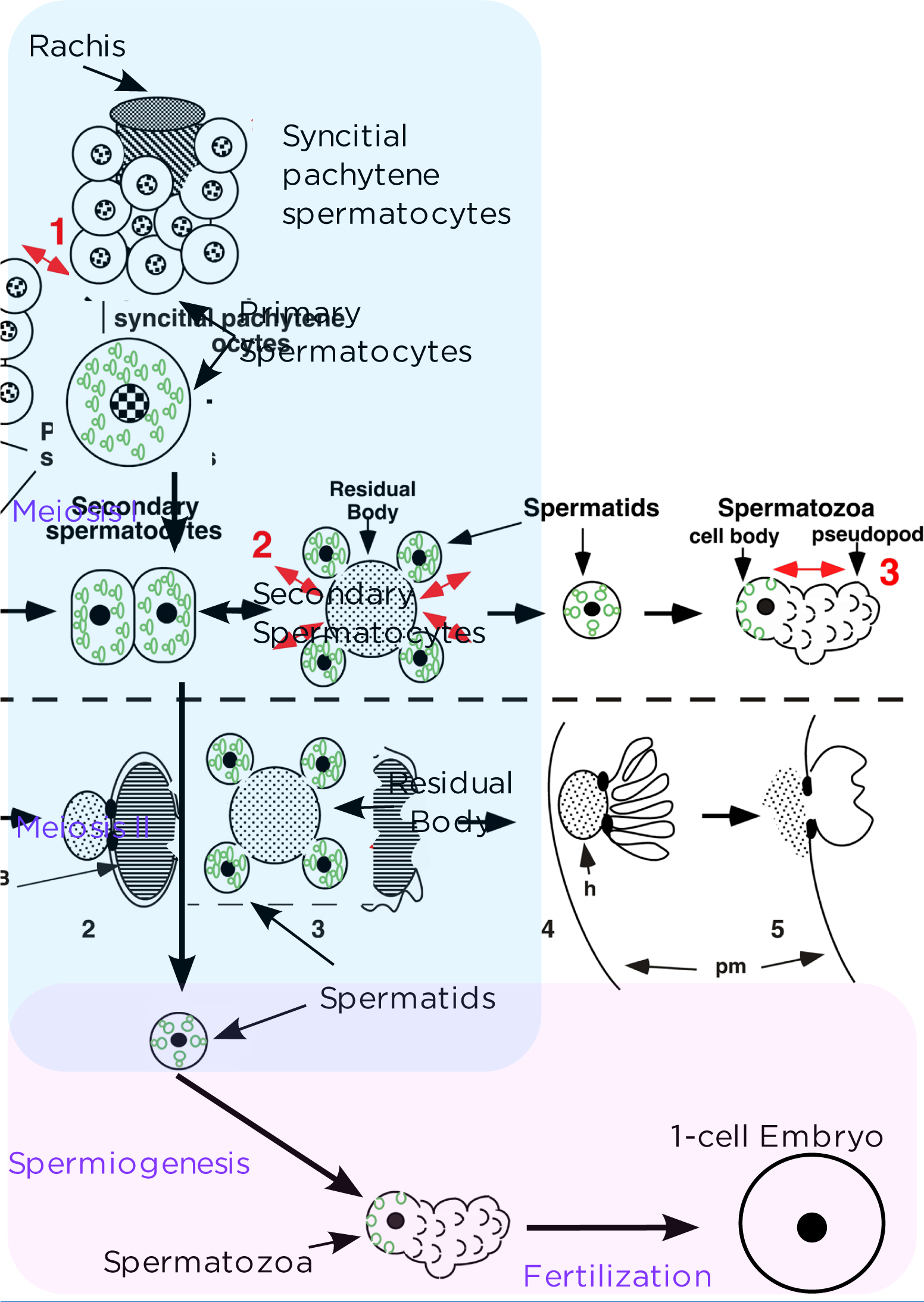
Spermatogenesis in *C. elegans*. Cell icons and information are taken from Figure 1 in (L’Hernault, 2006). Green icons are the fibrous body-membranous organelles (FB-MOs) that get selectively packaged with the spermatids while ribosomes, tubulin, and actin get left in the residual body. Stages shown in blue occur in the male and are expected to be found in the male atlas, while stages shown in pink occur in the female and are expected to be found in the female atlas.

In the dioecious *H. bakeri*, spermatogenesis only occurs in the male, but adults at 10 days post infection have already been mating, as evidenced by detection of eggs in host feces by 240 hours post-infection (Bryant, 1973). Therefore, sperm cells at various stages of maturation may be detected in both the male and female single-cell atlases. Assuming a similar process in *H. bakeri* as *C. elegans*, primary and secondary spermatocytes would be expected in the male atlas.

Spermatids would be expected in both the male and female atlases, while spermatozoa would be expected in the female atlas. If the undifferentiated spermatogonia form a syncytium, depending on their size, they may not be recovered well after filtration during processing or through the microfluidic channels of the Chromium system and so may not be detected in the male atlas. 10X Genomics lists the maximum tested cell size as 30 μm, though the channels are 70 μm in diameter (https://kb.10xgenomics.com/hc/en-us/articles/218170543-What-is-the-range-of-compatible-cell-sizes-).

Upon examining the marker genes for all clusters in the male and female atlases, annotated transcripts for major sperm proteins were found to be abundant cluster markers for male clusters 6, 12, 13, and 14, and for female clusters 7, 14, and 17 (Table S1 and S2). Moreover, there is high overlap in the major sperm protein cluster markers between male cluster 12, male cluster 6, male cluster 13, female cluster 14, and female cluster 7 (Table 3 and Figure 4). Additionally, annotated transcripts for ribosomal proteins, which were generally common cluster markers, were noticeably lacking as cluster markers for male clusters 6, 12, 13, 14, and 16, and for female clusters 4, 7, 13, 14, 15, and 17 (Table 3). Interestingly, the sole ribosomal protein cluster marker for male cluster 12 is the same feature as the sole ribosomal protein cluster marker for female cluster 17, while the sole ribosomal protein cluster marker for male cluster 13 is one of the two ribosomal protein cluster markers for female cluster 14 (Table S1 and S2). Transcripts annotated as tubulin or actin are not common enough as cluster markers to help distinguish potential sperm-related clusters from other clusters. Though the male and female clusters mentioned above are depleted in markers annotated to be tubulin or actin (Table S1 and S2).

**Figure 4.**
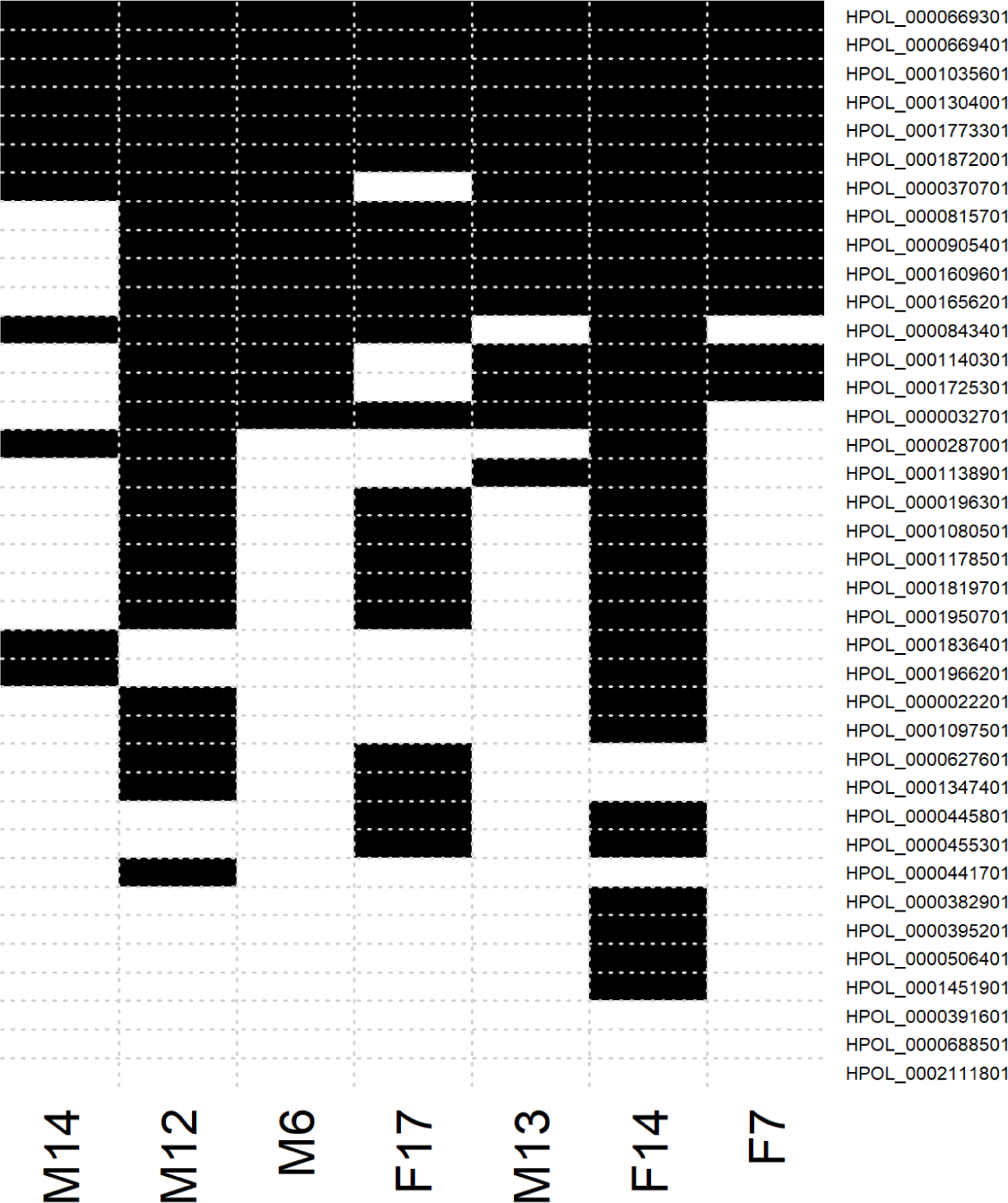
Distribution of major sperm protein cluster markers in putative sperm clusters. All genes from the *H. bakeri* genome annotated as major sperm protein are shown as rows. Black cells denote the presence of the gene (labels on the right) as a cluster marker for the cluster (labels on the bottom). White cells denote the absence of the gene as a cluster marker.

**Table 3.** Frequency of cell-type-informative cluster markers.

| (Atlas)<br>Cluster<br>Number | Number<br>of cluster<br>markers | Number<br>of<br>unique<br>cluster<br>markers | Number<br>of major<br>sperm<br>protein<br>markers | Number of<br>ribosomal<br>protein<br>markers | Number<br>of chitin-related<br>markers | Number of<br>chondroitin<br>proteoglycan<br>markers | Number<br>of<br>cuticle<br>collagen<br>markers |
| --- | --- | --- | --- | --- | --- | --- | --- |
| M0 | 260 | 36 | 0 | 68 | 0 | 0 | 0 |
| M1 | 368 | 22 | 0 | 23 | 0 | 0 | 0 |
| M2 | 237 | 5 | 1 | 70 | 0 | 0 | 0 |
| M3 | 252 | 46 | 0 | 22 | 0 | 0 | 0 |
| M4 | 259 | 72 | 0 | 9 | 0 | 0 | 0 |
| M5 | 323 | 12 | 1 | 83 | 0 | 0 | 0 |
| M6 | 241 | 0 | 15 | 0 | 0 | 0 | 0 |
| M7 | 582 | 52 | 1 | 22 | 0 | 0 | 0 |
| M8 | 602 | 41 | 3 | 10 | 0 | 0 | 0 |
| M9 | 152 | 21 | 0 | 63 | 2 | 5 | 0 |
| M10 | 496 | 34 | 0 | 23 | 0 | 0 | 0 |
| M11 | 426 | 73 | 0 | 86 | 0 | 0 | 0 |
| M12 | 1121 | 614 | 27 | 1 | 0 | 0 | 0 |
| M13 | 248 | 25 | 15 | 1 | 0 | 0 | 0 |
| M14 | 681 | 57 | 11 | 5 | 0 | 0 | 2 |
| M15 | 579 | 187 | 0 | 104 | 2 | 0 | 0 |
| M16 | 197 | 146 | 0 | 6 | 0 | 0 | 16 |
| F0 | 333 | 83 | 0 | 31 | 1 | 0 | 0 |
| F1 | 188 | 6 | 1 | 5 | 0 | 0 | 0 |
| F2 | 340 | 49 | 0 | 80 | 0 | 0 | 0 |
| F3 | 666 | 262 | 1 | 15 | 0 | 0 | 0 |
| F4 | 79 | 21 | 0 | 2 | 0 | 0 | 0 |
| F5 | 739 | 71 | 0 | 103 | 2 | 1 | 1 |
| F6 | 208 | 4 | 0 | 39 | 1 | 0 | 0 |
| F7 | 228 | 2 | 13 | 0 | 0 | 0 | 0 |
| F8 | 917 | 13 | 0 | 75 | 2 | 5 | 1 |
| F9 | 611 | 112 | 3 | 15 | 0 | 0 | 0 |
| F10 | 543 | 40 | 0 | 84 | 0 | 3 | 1 |
| F11 | 1107 | 50 | 0 | 49 | 2 | 8 | 2 |
| F12 | 306 | 44 | 0 | 15 | 0 | 0 | 0 |
| F13 | 142 | 6 | 0 | 2 | 2 | 8 | 0 |
| F14 | 1364 | 509 | 32 | 2 | 0 | 0 | 2 |
| F15 | 202 | 122 | 1 | 0 | 0 | 0 | 1 |
| F16 | 1093 | 253 | 1 | 25 | 2 | 8 | 1 |
| F17 | 771 | 107 | 21 | 1 | 0 | 0 | 0 |

I also examined the expression of orthologs of genes in *C. elegans* that may serve as useful sperm markers (Figure 5). In particular, *spe-6*, a casein I type serine threonine kinase that is important for FB-MO formation and also for spermatid maturation into spermatozoa (L’Hernault, 2006), and *spe-10*, an integral membrane protein which is required for proper FB-MO partitioning into the spermatids, both indicate male clusters 6, 12, and 13 and female clusters 7, 14, and 17 as containing cells with high expression of these transcripts.

**Figure 5.**
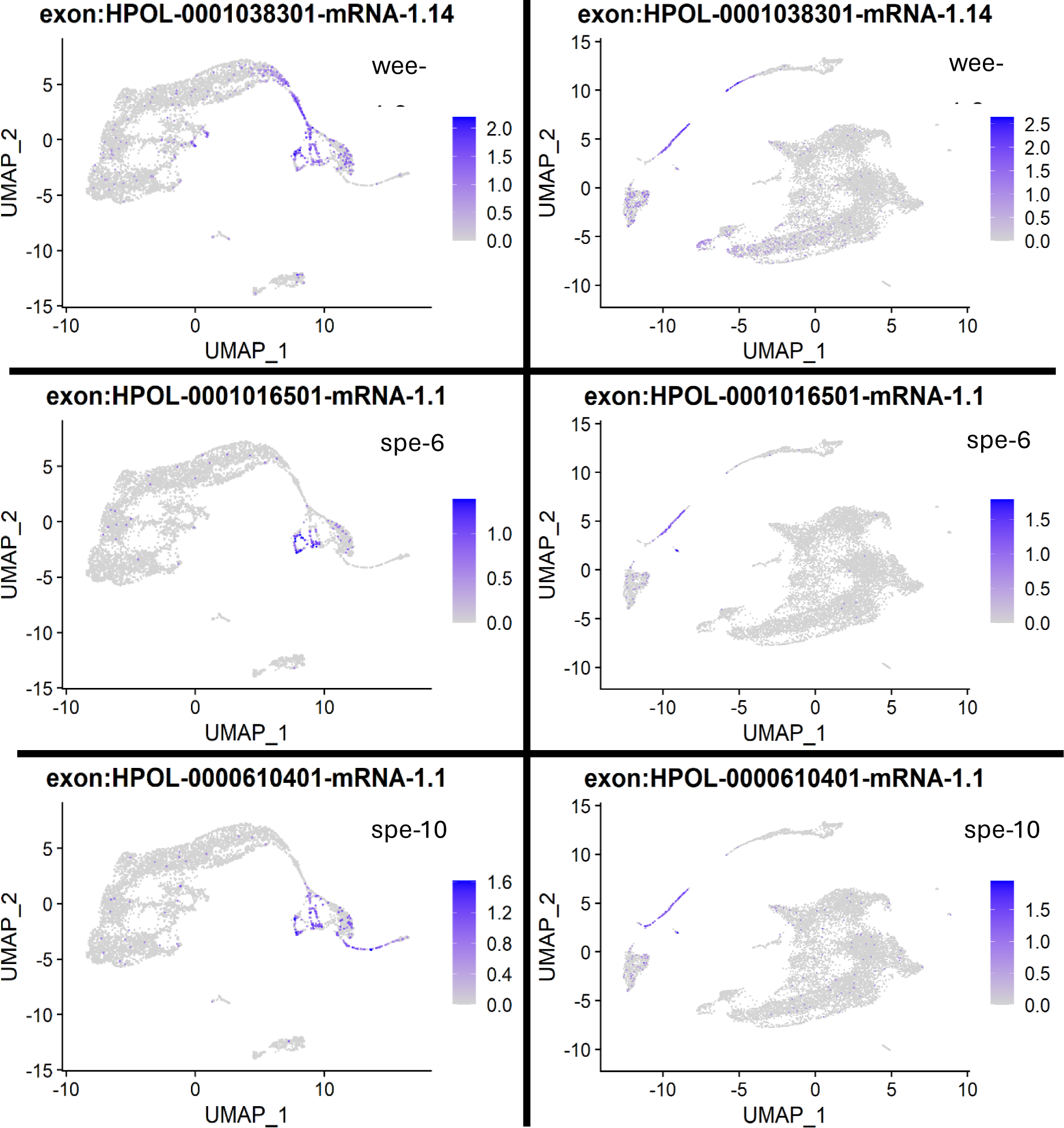
FeaturePlots of expression of orthologs of *C. elegans* genes important for spermatogenesis. Cells in the male (left) and female (right) atlases are coloured according to the expression level (SCT-normalized UMI count for that transcript for that cell) of the feature shown at the top of each plot. Genes shown include *wee-1.3*, a master regulator of cell divisions, especially important for spermatogenesis (L’Hernault, 2006), *spe-6*, a casein I type serine threonine kinase important for proper spermatid maturation (L’Hernault, 2006), and *spe-10*, an integral membrane protein required for proper formation of spermatids (L’Hernault, 2006).

LIGER cross-species analysis of *H. bakeri* and *C. elegans* scRNA-seq data (Table S3) clusters cells of male cluster 14 with either somatic gonad or intestinal cells of *C. elegans*, while *H. bakeri* cells of male clusters 12 and 6 cluster together with either germline or intestinal cell of *C. elegans*. Cells of male cluster 13 cluster with either germline or neural cells of *C. elegans*. *H. bakeri* cells of female cluster 7 cluster together with either germline or intestinal cells of *C. elegans*, while cells of female cluster 17 cluster together with germline cells of *C. elegans*, and cells of female cluster 14 cluster with germline, hypodermal, or neural cells of *C. elegans*.

Taken together, these results suggest that male clusters 14, 12, 6, and 13 are putatively sperm-related, with cluster 14 potentially being somatic gonad or spermatogonia at the beginning of spermatogenesis, cluster 12 potentially being primary spermatocytes, cluster 6 potentially being primary transitioning to secondary spermatocytes, and cluster 13 potentially being secondary spermatocytes transitioning into spermatids. Furthermore, the results suggest that female clusters 17, 14, and 7 are putatively sperm-related, with cluster 17 potentially being spermatocytes or immature spermatids (or potentially contamination of a male worm in a female sample), cluster 14 potentially being spermatids, and cluster 7 potentially being spermatozoa.

#### Annotating clusters: putative oocytes and eggs (and/or embryos)

In *C. elegans*, oocytes are fertilized by a spermatozoon in the spermatheca structure (L’Hernault, 2006). The now one-cell embryo (zygote) begins to form an egg, moves into the uterus, and develops to the roughly 30-cell stage over ∼150 minutes before the egg is laid (Altun & Hall, 2009a). During this time, the six-layer eggshell forms, cells divide and begin to differentiate, and a switch occurs from parental control of gene expression and cell patterning to zygotic control (the parental to zygotic transition or PZT) (Baugh et al., 2003; Stein & Golden, 2018). A summary of major events during this time and of the layers of the eggshell is given in Figure 6.

**Figure 6.**
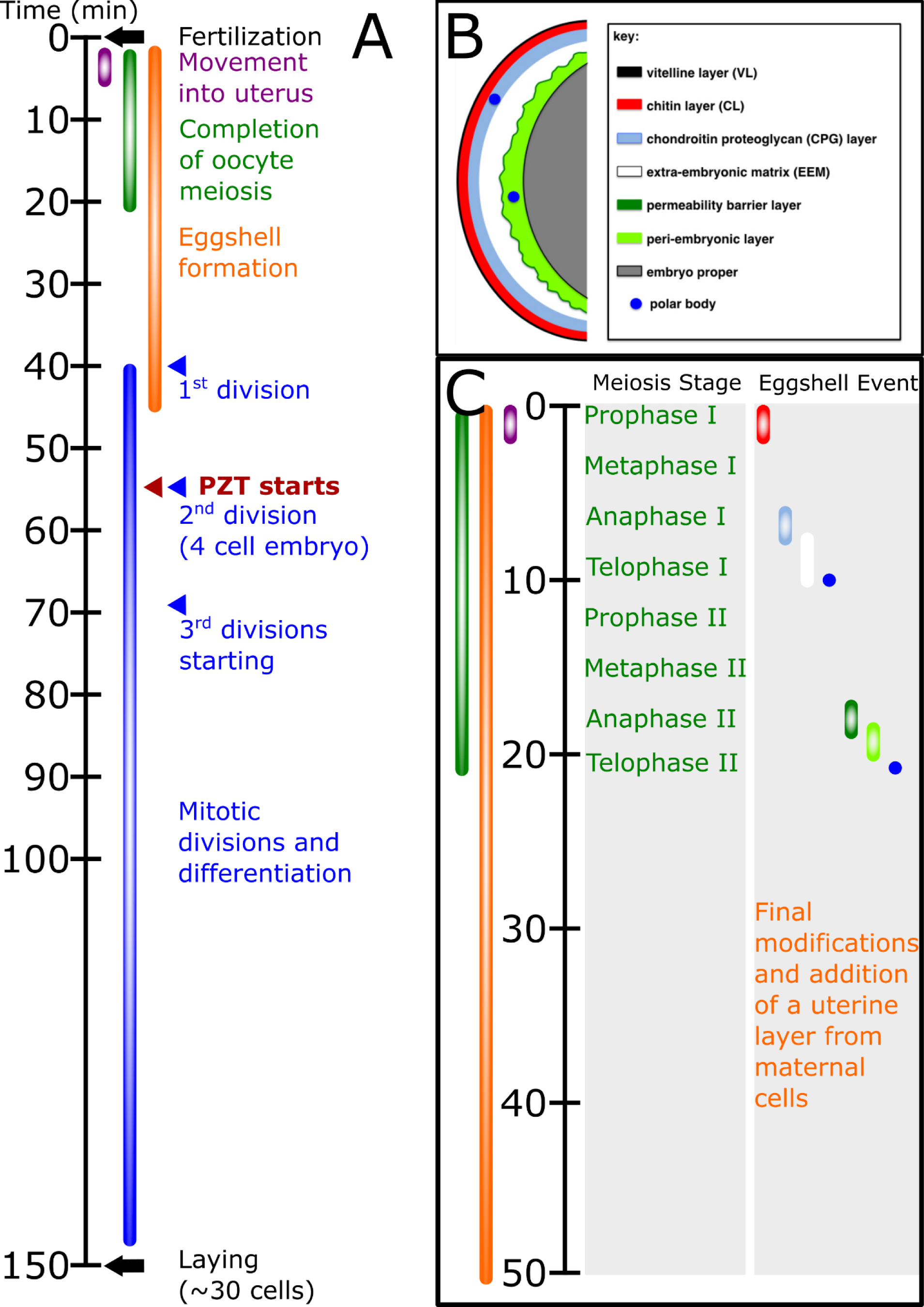
Timeline of early embryogenesis and egg formation in *C. elegans*. A) Major events in overall egg formation in *C. elegans* starting in the spermatheca structure of the mother. Coloured bars denote the approximate time of occurrence of the process described in the same colour to the right. PZT denotes the start of the parental to zygotic transition of control of gene expression. B) The layers of the *C. elegans* eggshell. Image is from WormBook, Figure 1 in (Stein & Golden, 2018). C) Timeline of the first 50 minutes to show the order and timing of the creation of the eggshell layers. Oocytes begin already having the vitelline layer. Coloured bars correspond to the layers as described in the legend of panel B. All information for this figure is taken from WormBook (Stein & Golden, 2018), with additional details from (Olson et al., 2012).

Assuming a similar process occurs in *H. bakeri*, the adult female is expected to contain: 1) adult female somatic cells, 2) oocytes in various stages of differentiation, and 3) embryos inside eggs up to the point of laying (and potentially shortly after laying depending on sample processing time). Since the eggshell has a different chemical composition to the adult cuticle, careful sample preparation and cell dissociation conditions could exclude the embryonic cells from a single-cell suspension by keeping them together within the intact egg. However, if the eggs are dissociated along with the adult females, embryonic cells, being small and round, should be recovered well in the 10X Genomics Chromium. Notably, all transcripts initially present in the one-cell embryo are of parental origin and thus the very early embryo would be expected to share many transcriptional features with parental cells, including, but not necessarily limited to, the gametes. Thus, embryonic cells from before the PZT may be indistinguishable from other cell types expected to be in the atlas, whereas embryonic cells from during or after the PZT should have distinct transcriptional profiles that could enable their identification.

Annotated transcripts for chitin-related terms (ex. Chitinase, chitin binding domain) were found to be cluster markers for female clusters 0, 5, 6, 8, 11, 13, and 16 and for male clusters 9 and 15 (Table S1 and S2). Additionally, transcripts annotated as chondroitin proteoglycan (3 or 4) were found to be cluster markers for female clusters 5, 8, 10, 11, 13, and 16 and for male cluster 9 (Table S1 and S2). Moreover, the two chitin binding domain transcripts that are markers for male cluster 9 and male cluster 15 are also the chitin-related markers for female clusters 5, 8, 11, 13, and 16. The eight chondroitin proteoglycan markers for female cluster 16 are the same transcripts as the eight markers for female cluster 13 and 11. These eight transcripts include all five of the male cluster 9 chondroitin proteoglycan transcripts and all of the chondroitin proteoglycan markers for female clusters 10, 8, and 5 (3, 5, and 1 markers, respectively). Given that chitin and chondroitin proteoglycan are important layers of the *C. elegans* eggshell (Figure 6B), but that eggshell formation starts before the PZT and thus relies at least in part on transcripts of parental origin, these clusters are potential candidates for being embryonic cells and/or oocytes. The similarity between male cluster 9 and the above female clusters suggests male cluster 9 may represent eggs contaminating the atlas.

The gene *rme-2* is the yolk receptor, expressed on oocytes and early embryos, responsible for the receptor-mediated endocytosis of yolk into unfertilized oocytes from the adult hermaphrodite intestine (Perez & Lehner, 2019). Expression of the ortholog of this gene in *H. bakeri* (HPOL_0000434101) implicates female clusters 2, 4, 5, 8, 10, 11, 13, and 16 and male cluster 9 as potentially oocytes and/or early embryonic cells (Figure 7). To attempt to identify the oocyte and early embryo clusters separately and resolve progeny cell clusters versus maternal cell clusters in the adult female atlas, we leveraged orthologs of collections of potential marker genes in *C. elegans* using the module scoring method in Seurat. This method calculates the average expression level of the set of genes provided minus the expression of a randomly selected set of control genes to yield a score, where higher positive values indicate cells with stronger expression of the genes provided. Firstly, genes to identify embryos from adult cells were selected based on the data from (Boeck et al., 2016). Specifically, we compared their bulk RNA-seq data on synchronized embryos (every 30 minutes starting at the four-cell stage) to adult sets and/or pre-PZT and/or pre-laying sets using the online portal the authors created (GExplore). The modules of genes in *C. elegans* resulting from these comparisons are in Table S4 and the *H. bakeri* ortholog module scores are plotted in Figure 8. In particular, the maternally enriched set of genes (matenr) represents expression in new embryos that corresponds to the time when they are still in the mother worm. Later timepoints (122 min or more) represent embryonic stages that are post-laying but pre-hatching. Secondly, genes identified by Warner and colleagues as broadly expressed and decreasing throughout their timepoints (every 90 minutes starting at egg laying) were used to distinguish embryos from adult cells (Warner et al., 2019). The module scores plotted on atlas UMAPs are shown in Figure 9 for Warner gene clusters 11, 18, and 20, which all consist of genes that showed particularly strong expression at the egg laying timepoint with a rapid decrease in expression at subsequent timepoints.

**Figure 7.**
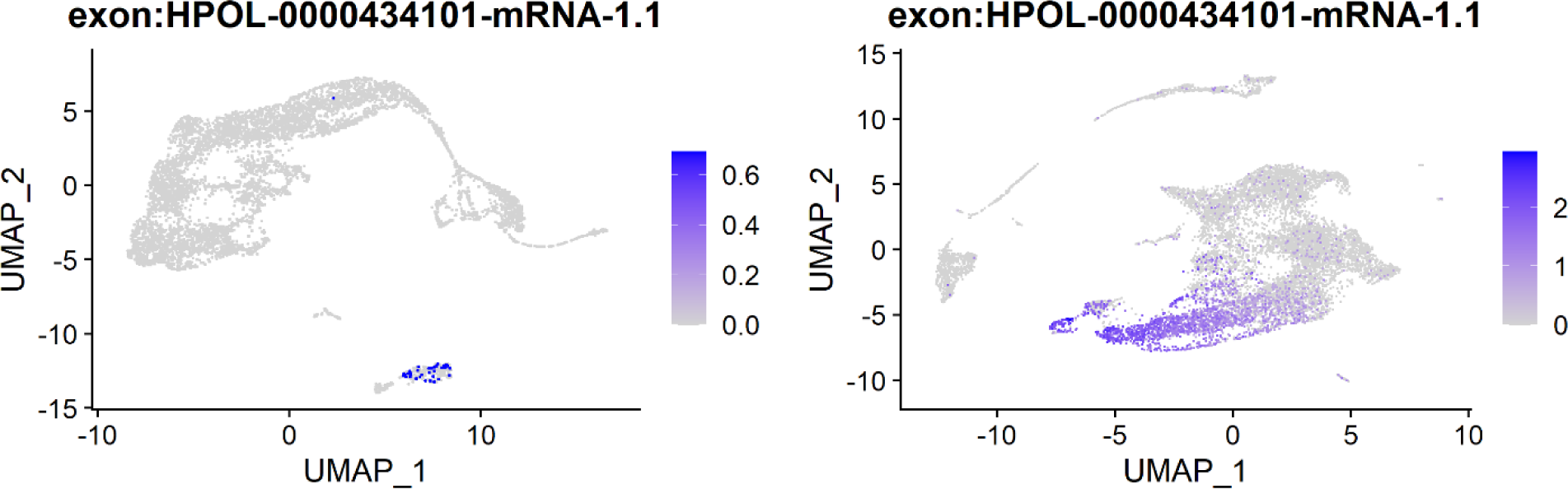
FeaturePlots of expression of the ortholog of *rme-2*. Cells in the male (left) and female (right) atlases are coloured according to expression level (SCT-normalized UMI count for that transcript for that cell).

**Figure 8.**
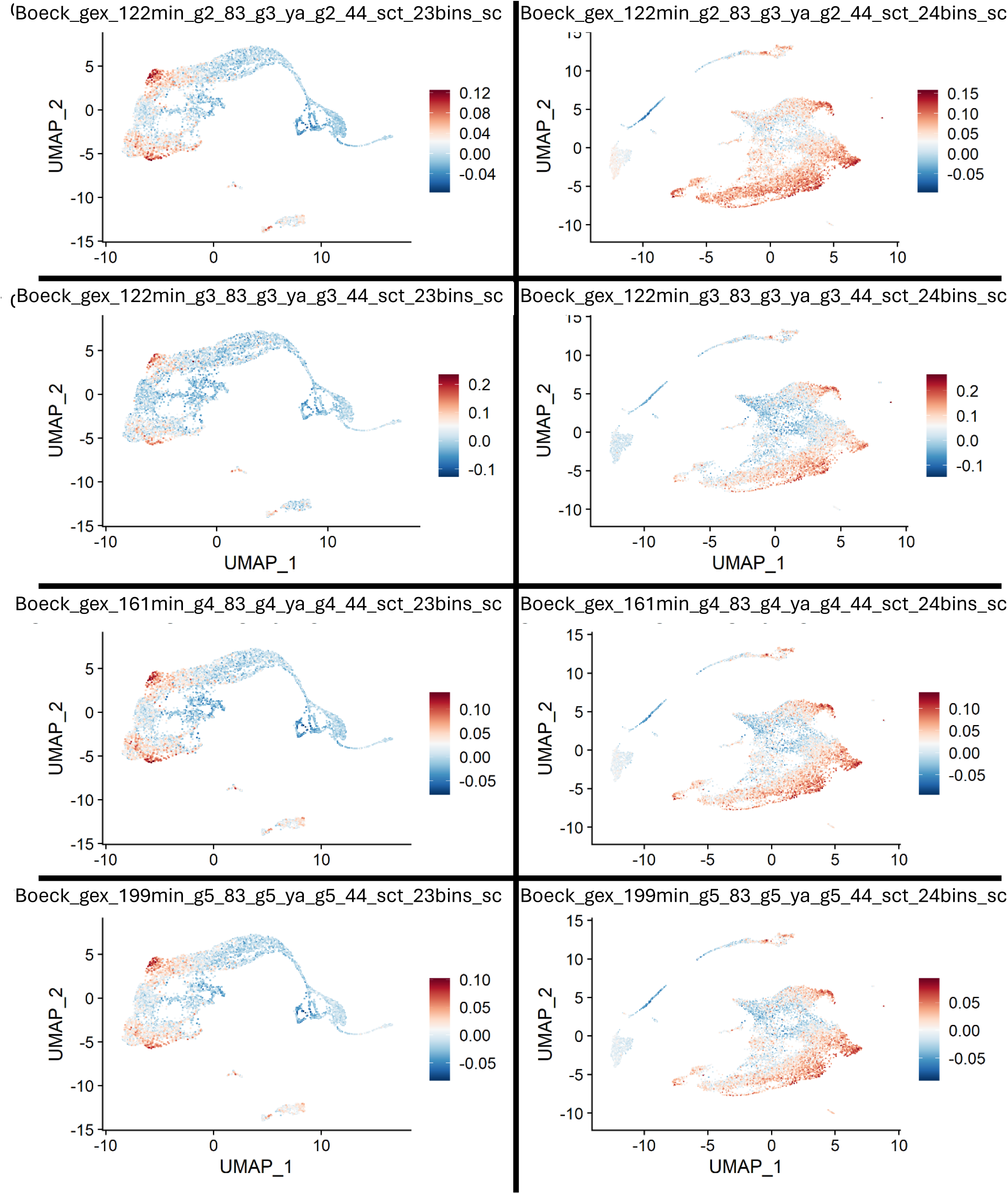
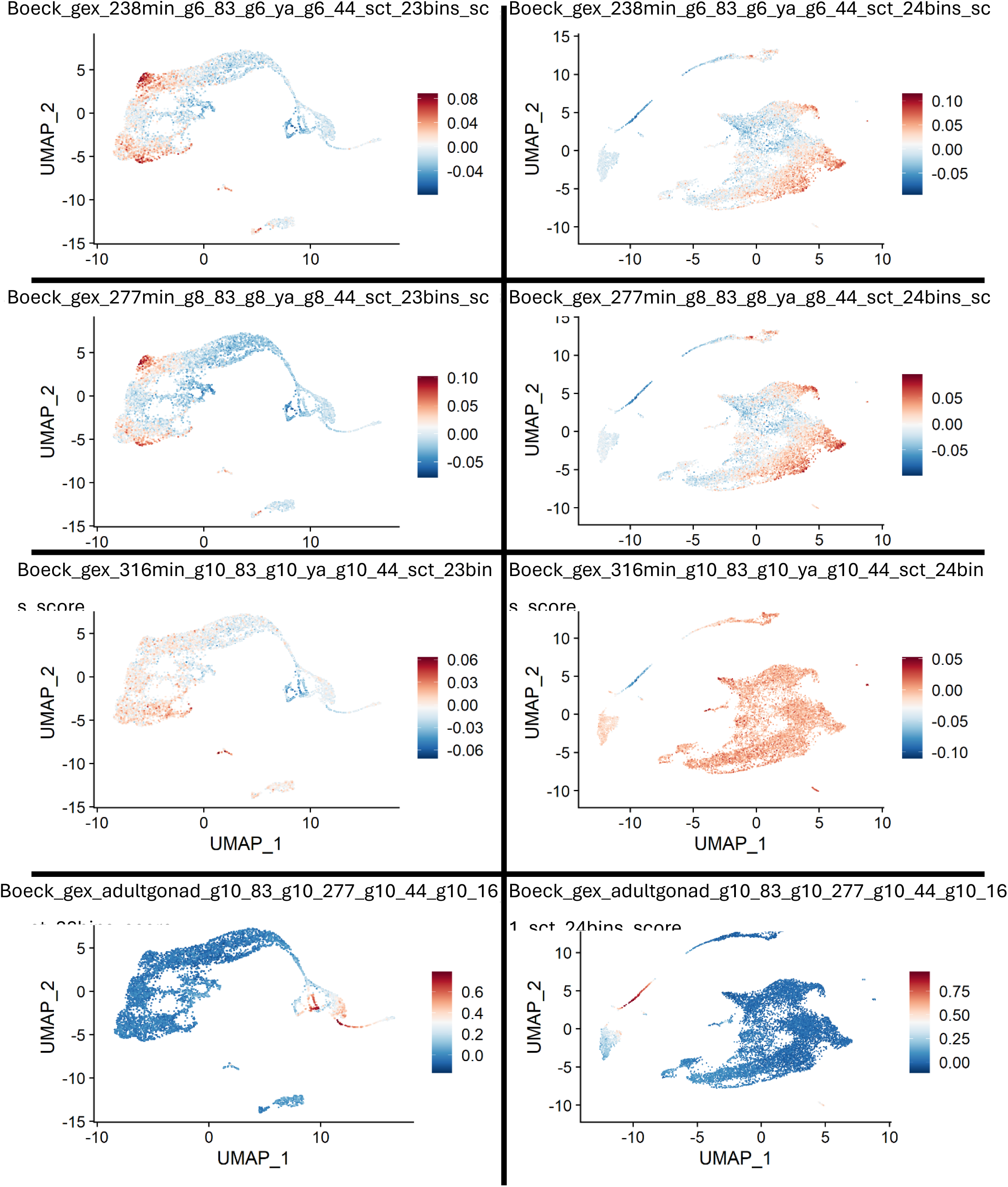
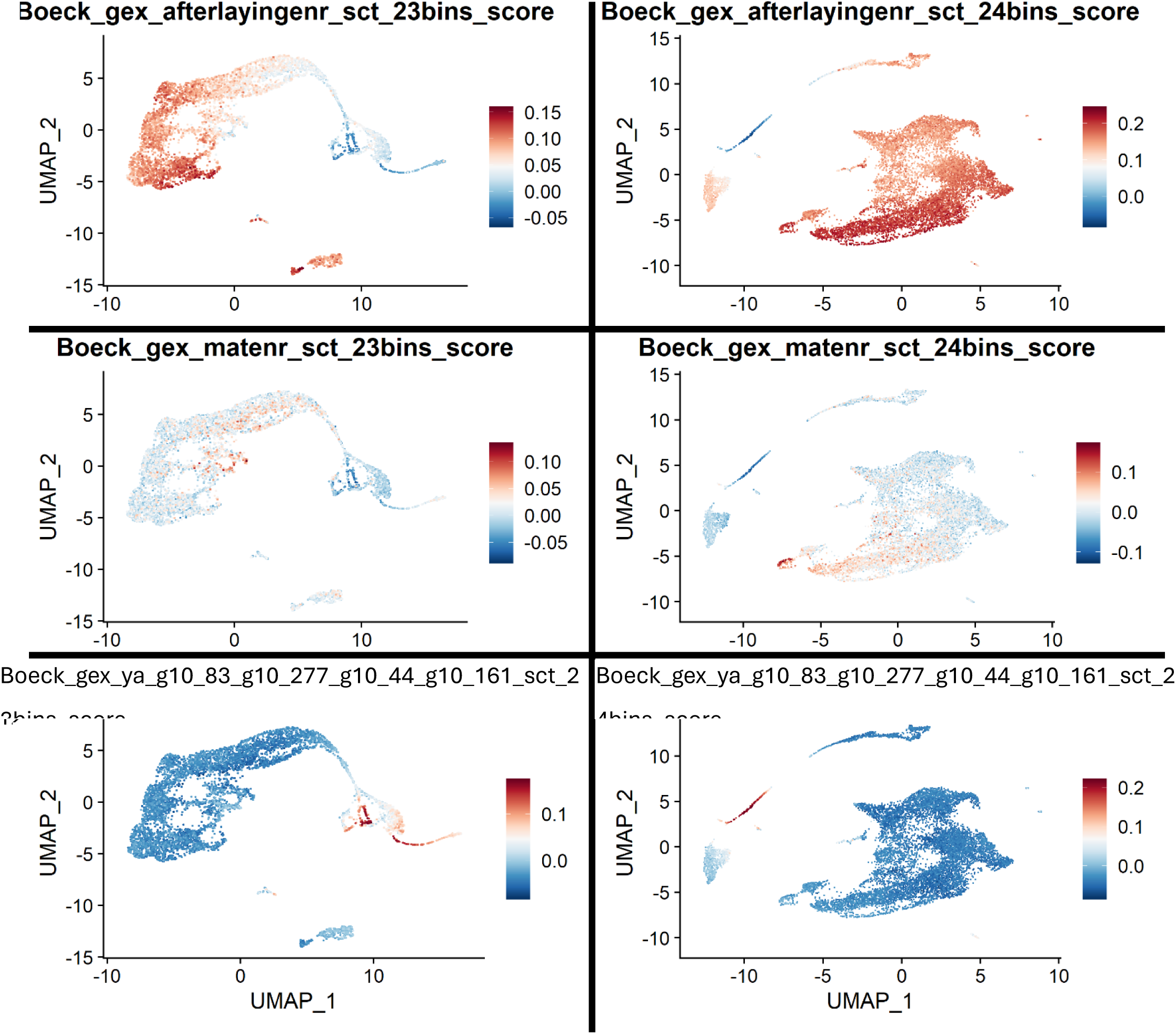
Boeck module scores to identify potentially embryonic cells. Cells are shown in either the male (left) or female (right) atlas, coloured according to their module score (See Section ‘Marker genes, C. elegans orthologs, and putative cluster annotation’) in the modules calculated from comparisons of the data from (Boeck et al., 2016) (See Section ‘Sources of marker gene modules’ for description of study). The full set of genes in each module is listed in Table S4, along with the details of the comparisons to get each of the modules. Module scores are relative and do not facilitate comparison to other modules or determination of a threshold score.

**Figure 9.**
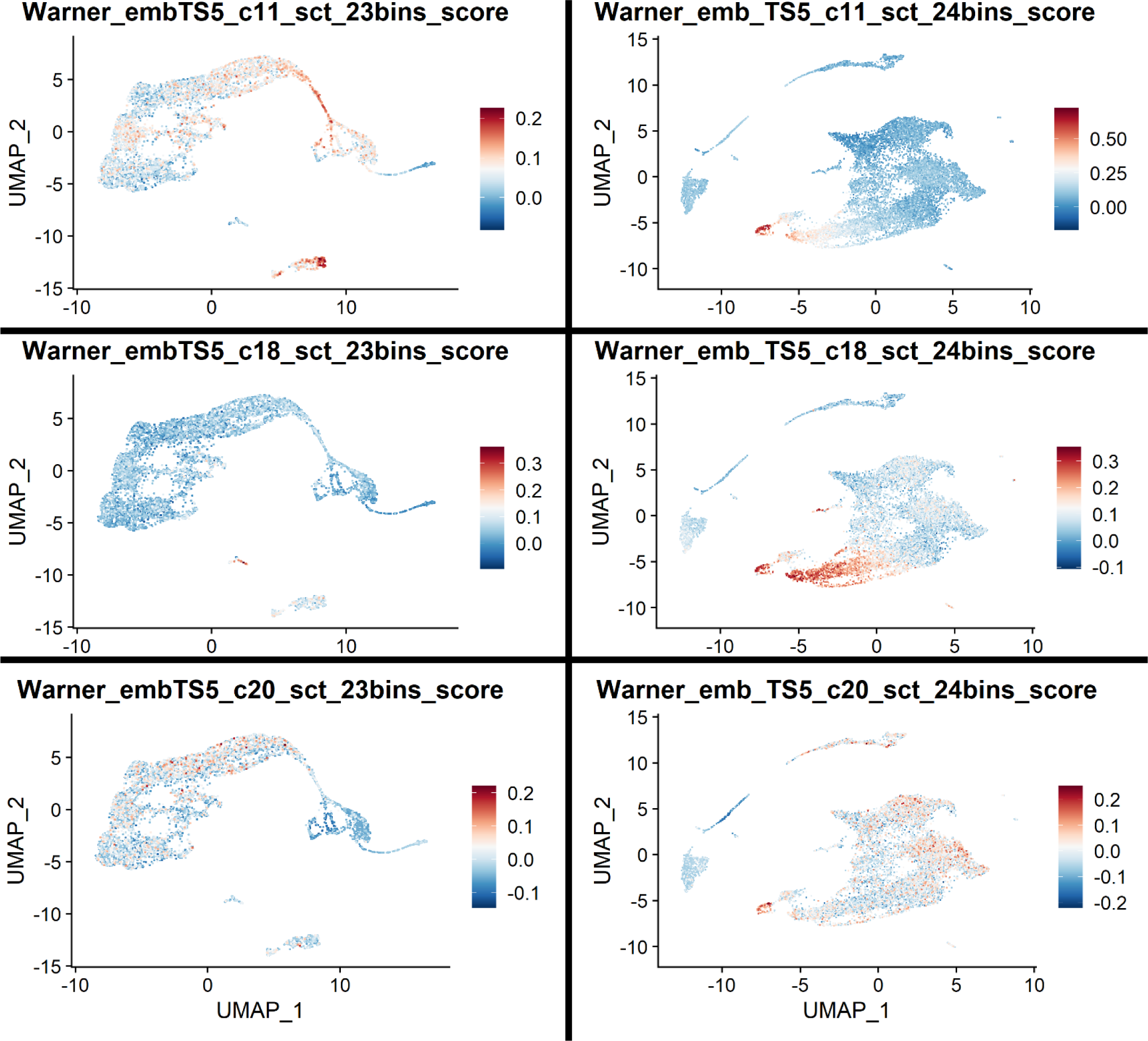
Warner module scores to identify potentially embryonic cells. Cells are shown in either the male (left) or female (right) atlas, coloured according to their module score (See Section ‘Marker genes, C. elegans orthologs, and putative cluster annotation’) in the modules calculated from sets of genes found to be highly expressed around the time of egg laying in (Warner et al., 2019) (See Section ‘Sources of marker gene modules’ for description of study). Module scores are relative and do not facilitate comparison to other modules or determination of a threshold score.

The LIGER cross-species analysis (Table S3) groups cells of female clusters 2, 5, 6, 8, 10, and 11 with germline cells of *C. elegans*, cells of female cluster 13 with either germline, pharynx, or neuron cells of *C. elegans*, and cells of female cluster 16 with either neuron, pharynx, germline, or support cells of *C. elegans*. Cells of male cluster 9 cluster with either germline, pharynx, neuron, or intestine cells of *C. elegans*.

Taken together, these results suggest that female clusters 2, 5, 6, 8, 10, 11, 13, and 16 are putatively oocyte- and/or embryo-related, with clusters 13, 16, and 11 potentially being oocytes or newly fertilized 1-cell embryos, cluster 8 potentially being young embryos before the PZT, cluster 10 potentially being embryonic, cluster 5 potentially being young embryos in eggs before laying, and clusters 2 and 6 potentially being young embryos in eggs near or shortly after laying. The results also suggest that male cluster 9 may be egg-related, which would suggest contamination of the ML and MP samples with either a few female worms or laid eggs present in the medium (Figure 10). The presence of so many cells in the female atlas (and any in the male atlas) whose expression profiles are consistent with oocytes and/or early embryos in eggs, regardless of their exact identity, confirms that the eggs were dissociated in the sample preparation procedures along with the adult worms. Separating the female atlas according to each cell’s library identity (Figure 10 and Table S5) reveals a previously unseen batch effect; a very high proportion of the cells in the clusters associated with oocyte/early embryo profiles come from the pronase library. Given the undefined nature of pronase (secretions of *Streptomyces griseus*), and the known depolymerization activity of pronase on chitosan (a deacetylated derivative of chitin) (Kumar, Gowda & Tharanathan, 2004), it is clear that the eggshells were dissolved during the pronase treatment, liberating the embryonic cells.

**Figure 10.**
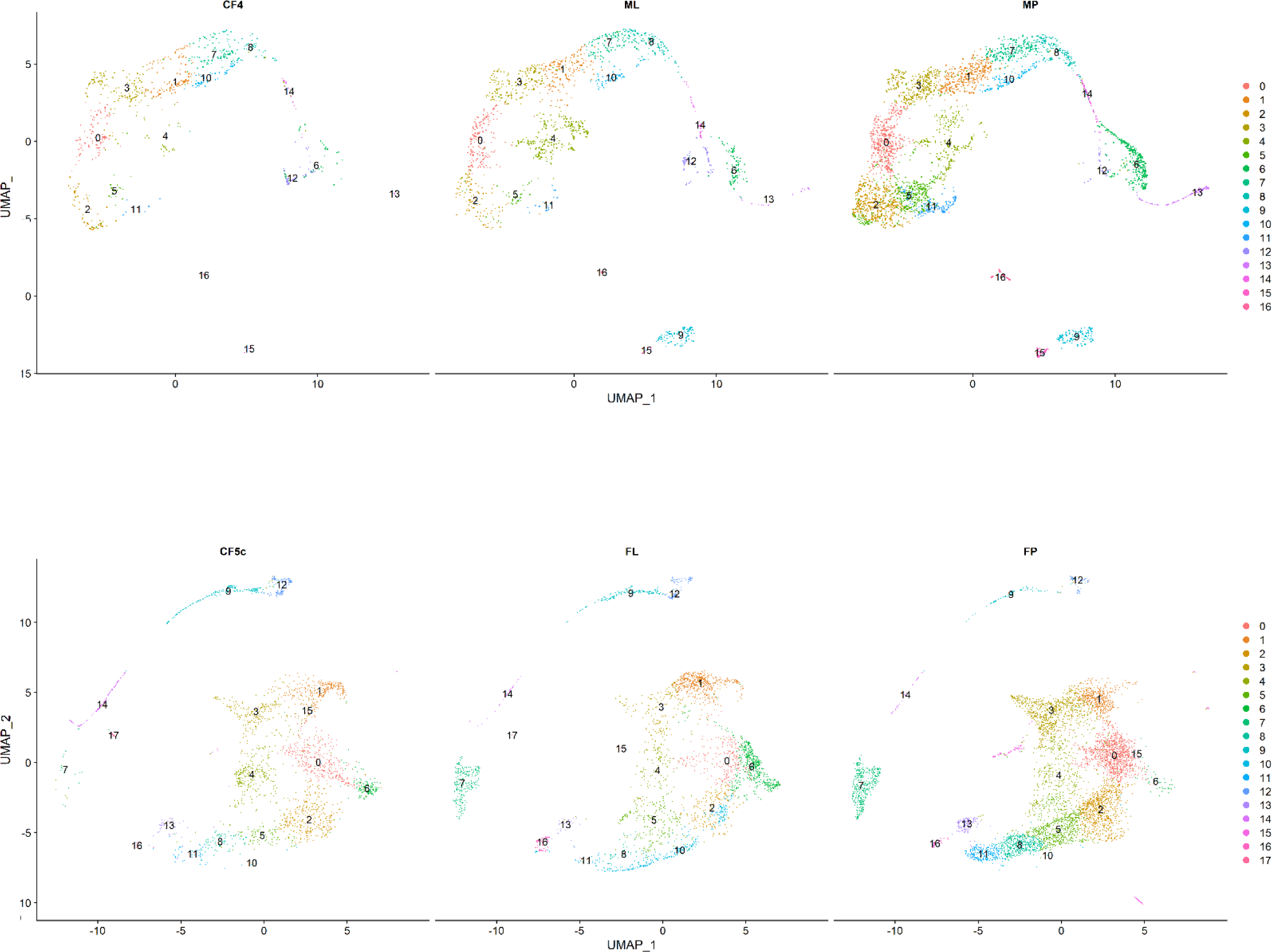
UMAPs of male (top) and female (bottom) atlases split by library. The title of each plot is the library name, with the third column in each panel representing the sample that was dissociated with pronase. Cells are coloured according to their cluster assignment. Note that cluster labels appear in the center point of all plotted cells from the cluster, which for female cluster 15 is a place that doesn’t actually include cells from the cluster.

#### Annotating clusters: putative hypodermis

In adult *C. elegans*, the outer epithelial layer of the worm, known as the hypodermis, consists of several large syncytia covering the main body and several smaller single nucleate cells at the head and tail (Altun & Hall, 2009b). External to the hypodermis is an exoskeleton layer, known as the cuticle, that is composed of collagen proteins, insoluble proteins called cuticlins, glycoproteins, and lipids (Page & Johnstone, 2007). During development, with each shedding of the cuticle (molting) the hypodermis synthesizes and secretes the components needed to build the new cuticle (Page & Johnstone, 2007). The over 170 cuticle collagens show temporal- and stage-specific expression patterns as the cuticle for each stage of development is different and has different composition (Page & Johnstone, 2007). Additionally, certain cuticle collagens continue to be expressed in adult hypodermis, even after cuticle synthesis, and are involved in maintaining the barrier function of the cuticle (Sandhu et al., 2021).

Assuming a similar structure to the hypodermis in *H. bakeri*, most of the hypodermis, by surface area, would exist in large multi-nucleate syncytia that may be too large to be recovered well after filtration or through the microfluidic channels of the 10X Genomics Chromium. However, the smaller hypodermal cells at the head and tail should be recovered. Moreover, as the only tissue involved in creating the cuticle, cuticle-related components should serve as markers of hypodermal cells.

Annotated transcripts for cuticle collagen were found to be cluster markers for male clusters 14 and 16 and for female clusters 5, 8, 10, 11, 14, 15, and 16 (Table S1 and S2). We also examined the expression of orthologs of known cuticle-related genes from *C. elegans*, including the cuticlin *cutl-18*, the tetraspanin *tsp-15*, the prolyl-4 hydroxylase *phy-2*, and the cuticle component *rol-1*, which when mutated causes a roller phenotype (Figure 11). Finally, we calculated module scores for the set of genes found to be unique to the hypodermis from bulk RNA-seq of hypodermal cells sorted from adults of a *pY37A1B.5::gfp* reporter strain (Kaletsky et al., 2018).

**Figure 11.**
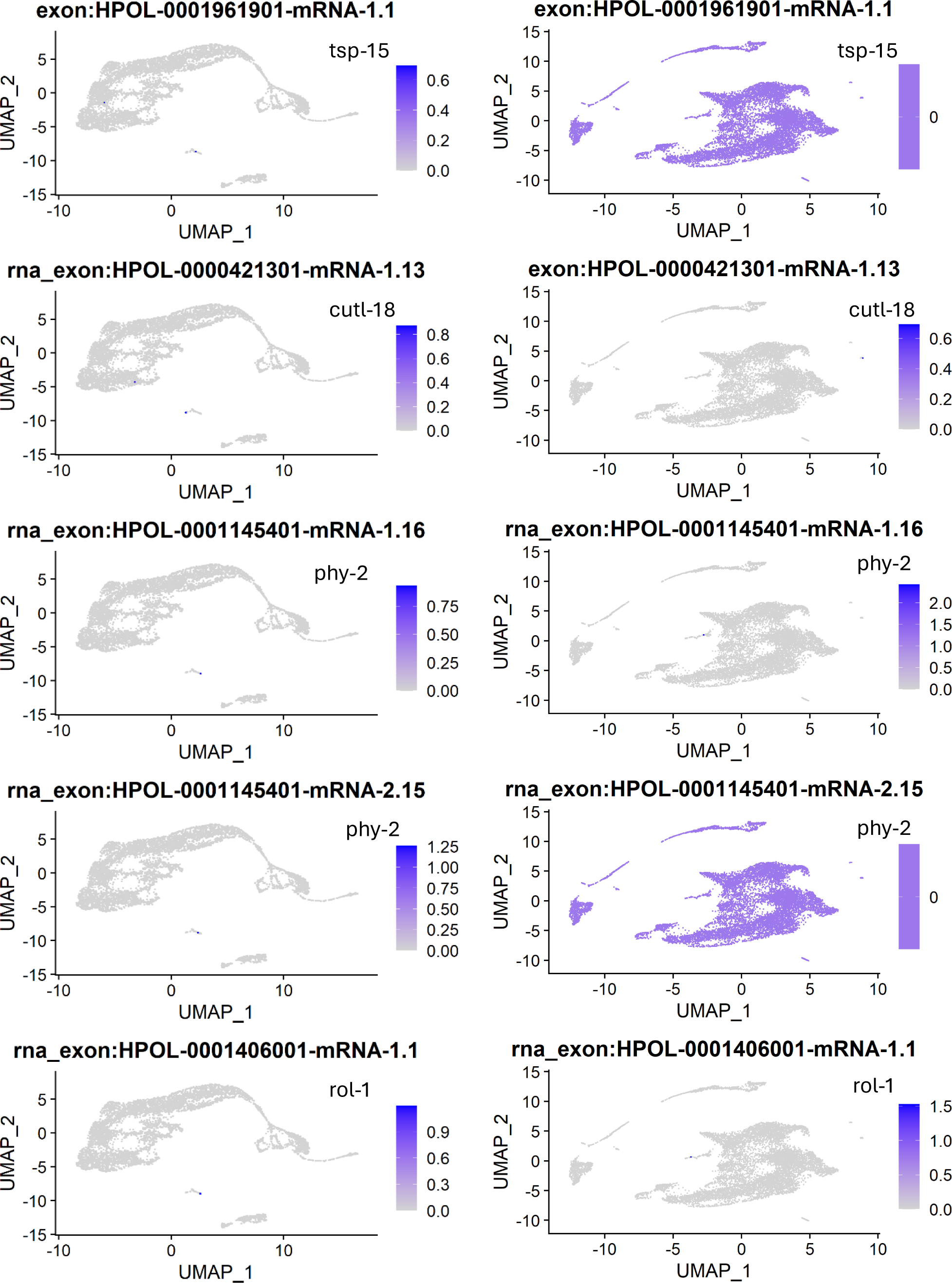
FeaturePlots of expression of orthologs of *C. elegans* genes involved in the cuticle. Cells in the male (left) and female (right) atlases are coloured according to the expression level (SCT-normalized UMI count for that transcript for that cell) of the feature shown at the top of each plot.

The LIGER cross-species analysis (Table S3) clusters cells of male cluster 16 with neuron, hypodermis, germline, somatic gonad, and seam cells (hypodermis) of *C. elegans*. Cells of female cluster 15 are clustered with body wall muscle, neurons, pharynx, germline, and somatic gonad cells of *C. elegans*.

Taken together, these results suggest that male cluster 16 and female cluster 15 putatively include adult hypodermis. Of note, female clusters 5, 8, 10, 11, and 16 are putatively associated with embryos, which would be synthesizing the first cuticle of the L1 worm that will hatch from the egg once fully developed. The lack of high scores in these clusters when using the Kaletsky adult unique hypodermal genes as a module likely reflects the significant changes in gene expression that occur as the hypodermis fully matures into an adult tissue from its embryonic precursors. Interestingly, the remaining clusters mentioned above, male cluster 14 and female cluster 14, are both putatively sperm-associated. While sperm themselves are not involved in cuticle synthesis and have no direct reason to have cuticle collagen transcripts, the idea that expression of certain components of the first cuticle (particularly various collagens) are under paternal control in a developing embryo is an intriguing possibility (See below section on embryogenesis).

#### Annotating clusters: putative intestine

The intestine is the largest tissue and accounts for roughly one third of the total cell volume in adult *C. elegans* (Froehlich, Rajewsky & Ewald, 2021). It is composed of 20 cells, in pairs, that have 32n nuclei (in contrast to most other tissues which are diploid) (McGhee, 2007). Additionally, some of the cells have two nuclei, such that the entire intestine can have 30–34 nuclei in total (McGhee, 2013). In addition to being the site of digestion and absorption, the intestine is also the major area for macromolecule storage (McGhee, 2007).

Assuming similar numbers, volume, and structure to the intestine in *H. bakeri* as in *C. elegans*, intestinal cells should be present in the atlases of both the male and female worms. However, the types of proteins characteristic of core intestinal functions (like digestive enzymes, proteases, lipases, or proteins involved in carbohydrate catabolism) are expressed broadly enough throughout the different cells of the worm that they will not serve as transcriptional markers to distinguish clusters of intestinal cells from non-intestinal cells. Rather, genes verified to be transcribed only in the intestine are required. One such key marker gene is *elt-2*, the master regulator of the intestine cell fate (McGhee, 2007), which has an ortholog in *H. bakeri* (HPOL_0001764901). However, this gene is only detected in a few cells and thus does not identify clusters of intestinal cells. Whether this is because this transcription factor does not itself need to be highly transcribed to perform its function (and thus is not detected well with this method), or because the *H. bakeri* ortholog does not have the same function as *elt-2* is unclear. The *Ascaris suum* ortholog of this gene was found to be highly expressed in bulk RNA-seq analysis of dissected intestine (Rosa, Jasmer & Mitreva, 2014). Of note, whole worm expression of this transcript in *H. bakeri* puts it among the top 51% of transcripts in worms of the same age as used here (Pollo et al., 2023).

I was able to find four sets of genes in *C. elegans* whose orthologs in *H. bakeri* may serve as useful modules to identify intestinal cells. The first set (Blazie_int_uniq) comes from (Blazie et al., 2015), who sequenced from mixed-stage worms RNA that was bound to a polyA-binding protein that was expressed under the control of an intestinal promoter (*ges-1*). Relative to the other tissues they examined (pharyngeal muscle and body wall muscle), they were able to define a set of 4091 genes uniquely expressed in the intestine. The second set comes from (Haenni et al., 2012), who sequenced 3’ ends of transcripts within intestinal nuclei of mixed-stage worms that were obtained from fluorescence-activated nuclei sorting. By comparing a sample of sorted intestinal nuclei to an unsorted sample, they were able to identify 2456 genes with higher expression in the intestinal nuclei. The third set comes from (Kaletsky et al., 2018), who sequenced from adult worms RNA from intestinal cells that were obtained by cell sorting of a *Pges-1::gfp* reporter strain. By comparing to other tissues (hypodermis, neurons, and muscle) they were able to define genes enriched in (highly expressed and significantly differentially expressed relative to the average expression of all the other tissues) or unique to (highly expressed and significantly differentially expressed relative to each of the other tissues) the intestine. The fourth set comes from a review of the literature, relying heavily on WormBook (McGhee, 2007). It is also worth noting that additional modules could be calculated in the future from re-analysis of the bulk RNA-seq datasets from dissected intestine from *A. suum* (Rosa, Jasmer & Mitreva, 2014) and *H. contortus* (Laing et al., 2013) and retrieval of the orthologs between these worms and *H. bakeri*.

The intestinal module scores plotted on the UMAPs of the male and female atlases are shown in Figure 12. Since the UMAP is a projection of highly multidimensional data into two-dimensional space, and the cluster assignment of a cell is not always obvious when comparing to a UMAP that is coloured by cluster (Figure 13), we opted to analyze the cluster assignments of the cells based on module score directly, rather than relying on visual inspection of the scores plotted on the UMAPs. Every cell is assigned a score, but the distribution of scores can vary wildly between modules (Figure 14), so a cutoff value to assign an ID to a cell based on module score is not appropriate. A quantile cutoff value may be more appropriate when there is confidence that a certain proportion of the cells recovered are the cell type reflected by the module (ex. If 10% of the atlas is intestine then the cells within the top 10% of intestine module scores are probably those intestinal cells). However, that is not the case here. We reasoned that if a random sample of the cells is taken and their cluster assignment checked, we should see the cell clusters get represented according to their overall frequency in the atlas. However, if we sample the cells in order of decreasing module score, we should see the main clusters containing the cells of interest get represented first. We can then calculate the fraction of each cluster that is represented when sampling different amounts of the atlas, remembering that only positive module scores would indicate an increased expression of the genes in the module and that linear increase in the fraction of a cluster represented would be the equivalent of random assignment of high scoring cells to cell clusters. Therefore, clusters whose fractional representation increases faster than linear (above the diagonal line) in positive module scores (to the left of the vertical line) are clusters that contain high scoring cells for that module at a frequency higher than expected by chance alone. The results of this for the intestinal modules are shown in Figure 15.

**Figure 12.**
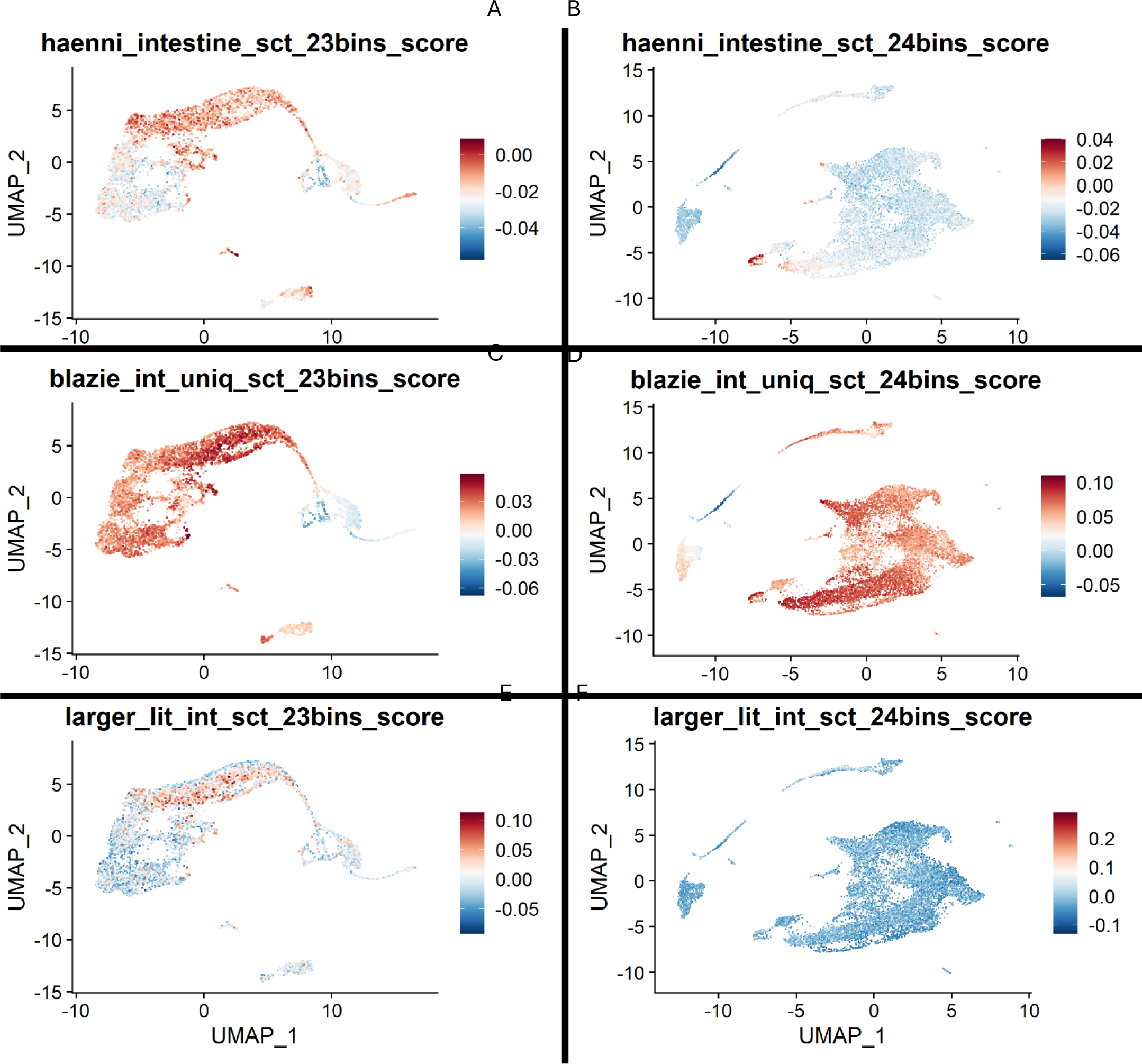
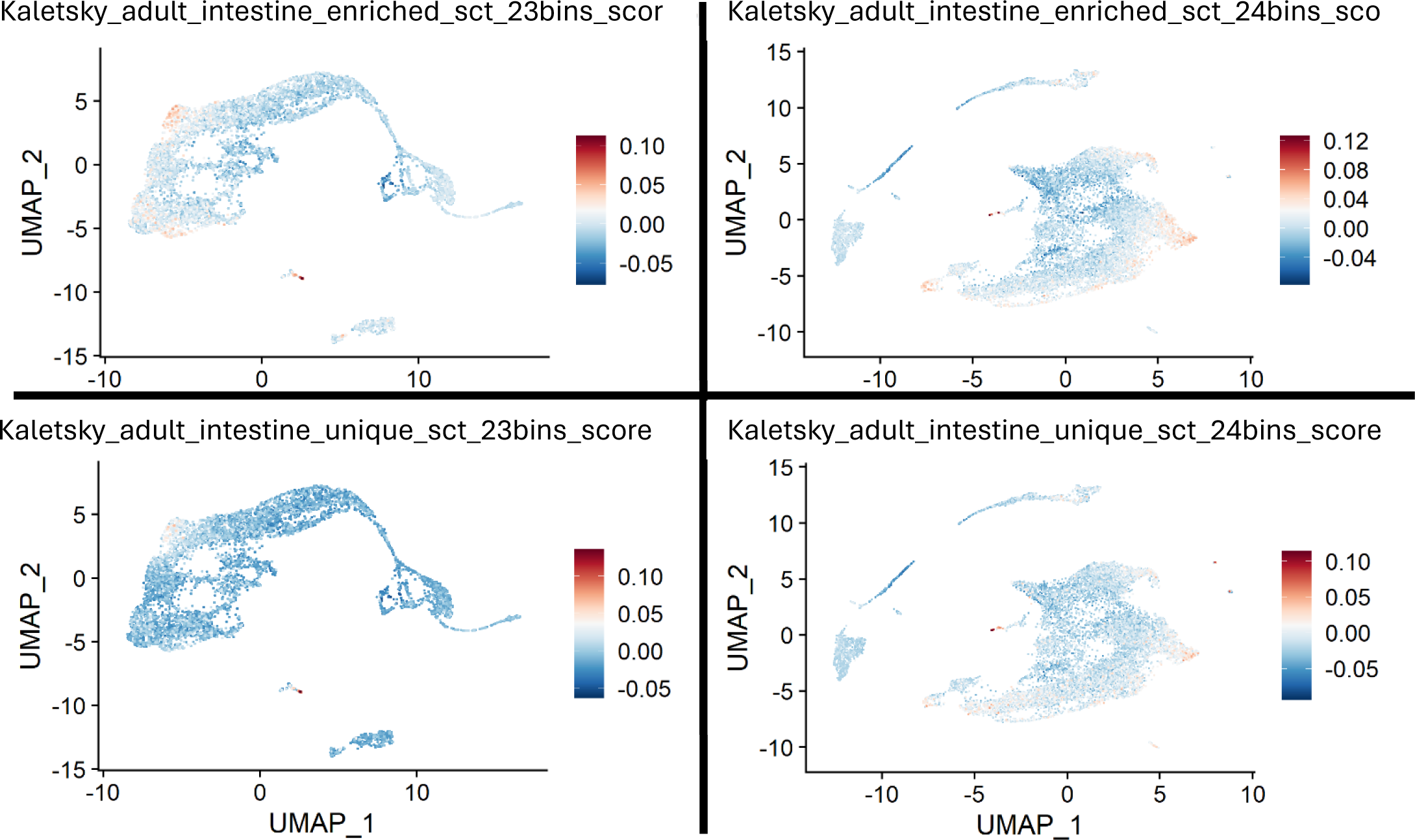
Intestinal module scores to identify potential intestinal cells. Cells are shown in either the male (left) or female (right) atlas, coloured according to their module score (See Section ‘Marker genes, C. elegans orthologs, and putative cluster annotation’) in the intestine-related modules calculated from different sets of genes: A) and B) genes found to be up-regulated in intestinal nuclei relative to unsorted nuclei in mixed-stage worms (Haenni et al., 2012), C) and D) genes found to be unique to intestinal cells relative to pharyngeal muscle and body wall muscle in mixed-stage worms (Blazie et al., 2015), E) and F) genes found to be important for the intestine in literature search and WormBook (McGhee, 2007), G) and H) genes found to be enriched in intestinal cells relative to hypodermis, neurons, and muscle in adult worms (Kaletsky et al., 2018), and I) and J) genes found to be unique to intestinal cells relative to hypodermis, neurons, and muscle in adult worms (Kaletsky et al., 2018). Module scores are relative and do not facilitate comparison to other modules or determination of a threshold score.

**Figure 13.**
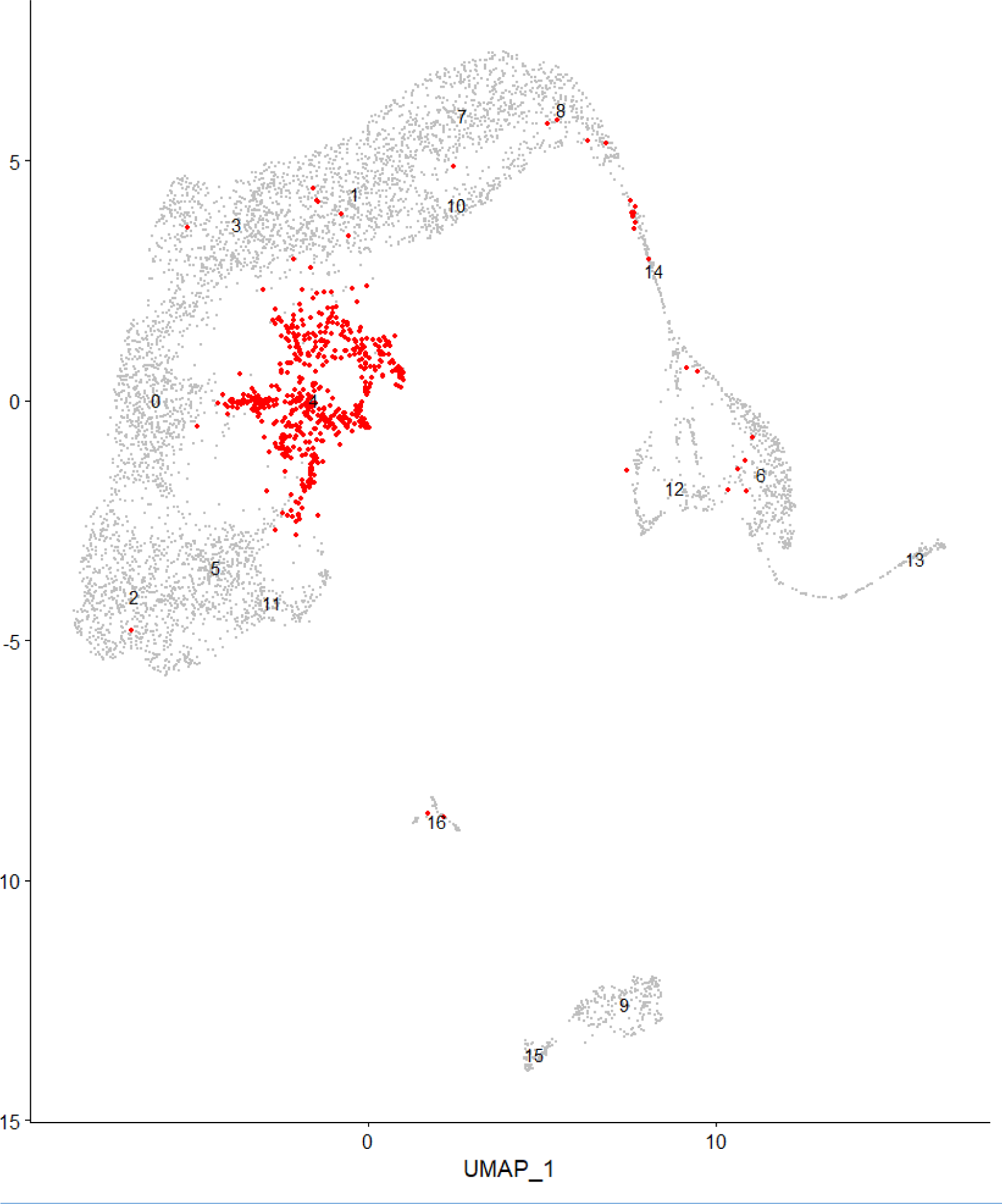
UMAP of the male atlas, highlighting cluster 4. Cells of the male atlas are shown in grey, except for the cells assigned to cluster 4, which are shown in red.

**Figure 14.**
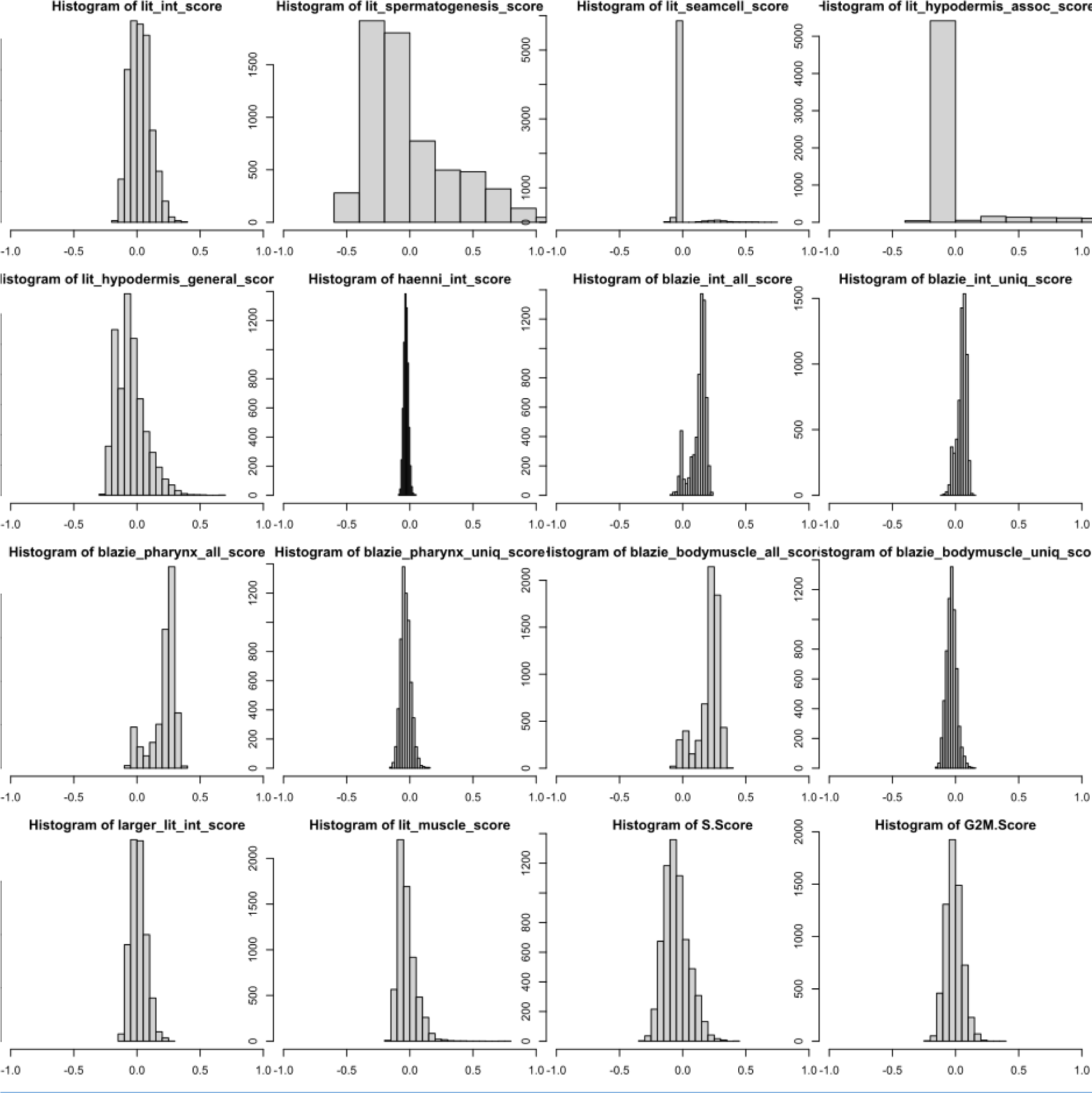
Distribution of scores of various modules. The distributions for a selection of 16 module scores (See Section ‘Marker genes, C. elegans orthologs, and putative cluster annotation’) are shown as histograms. The variability in range of values and skew of the overall distribution is highlighted.

**Figure 15.**
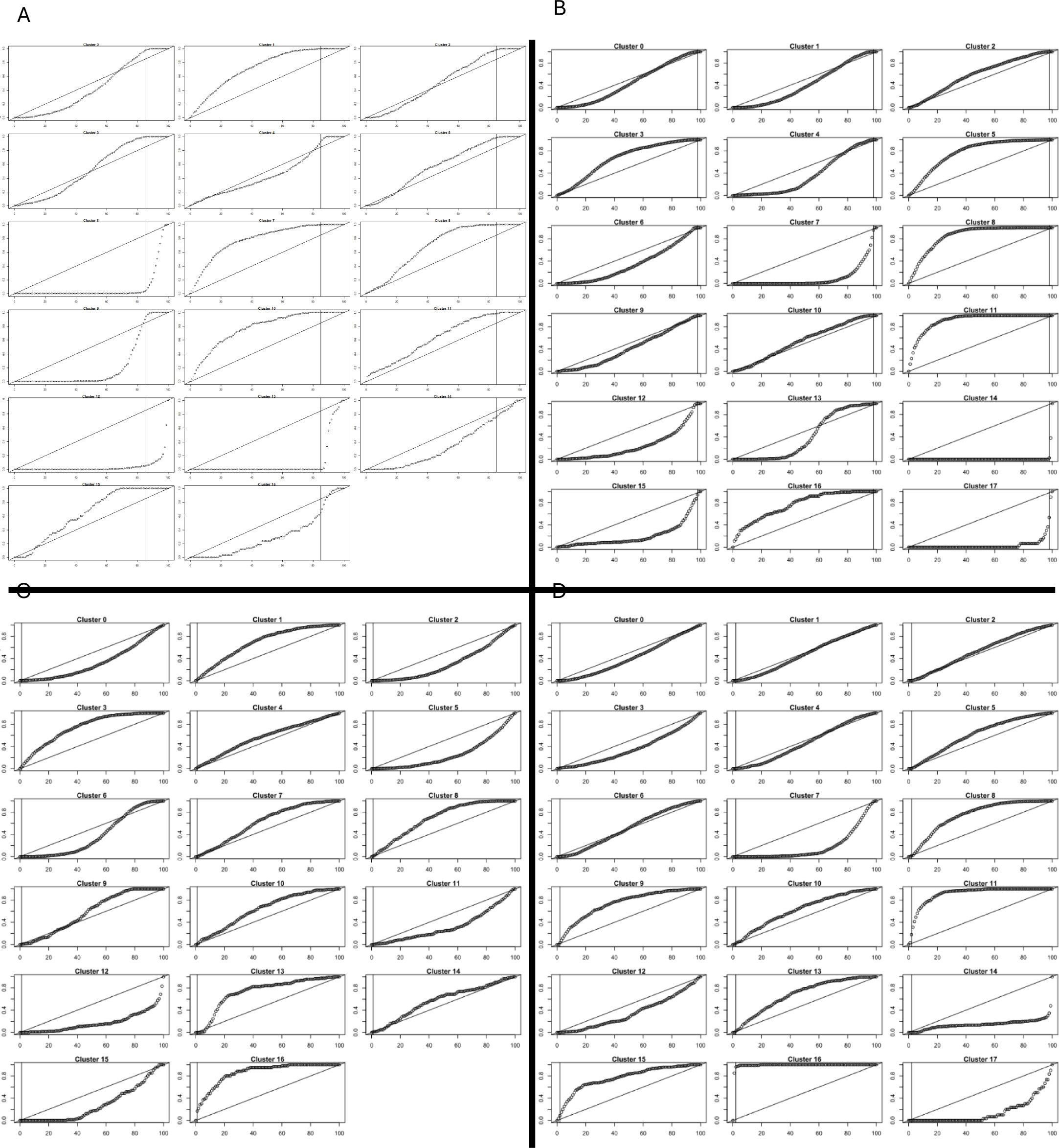
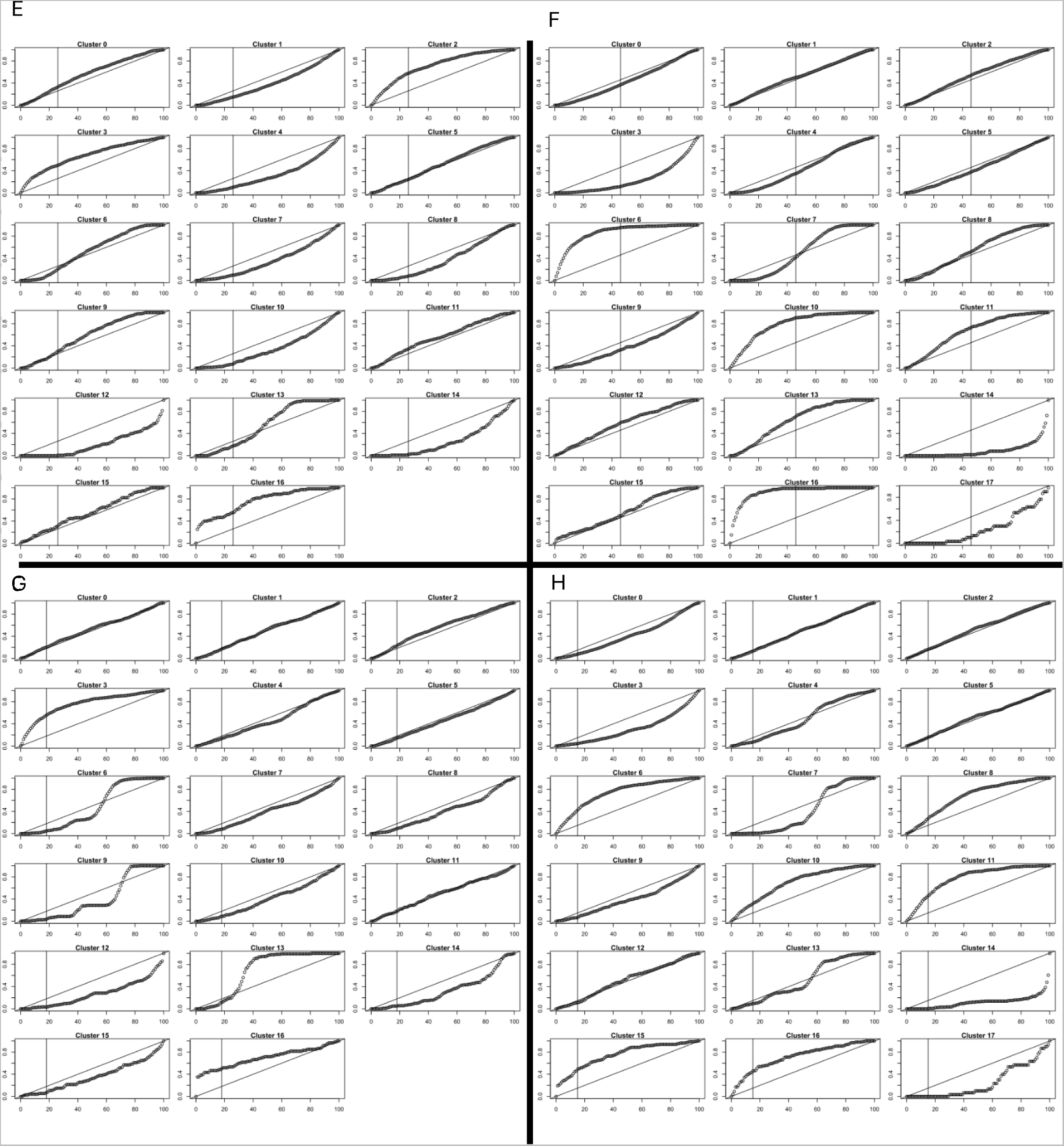
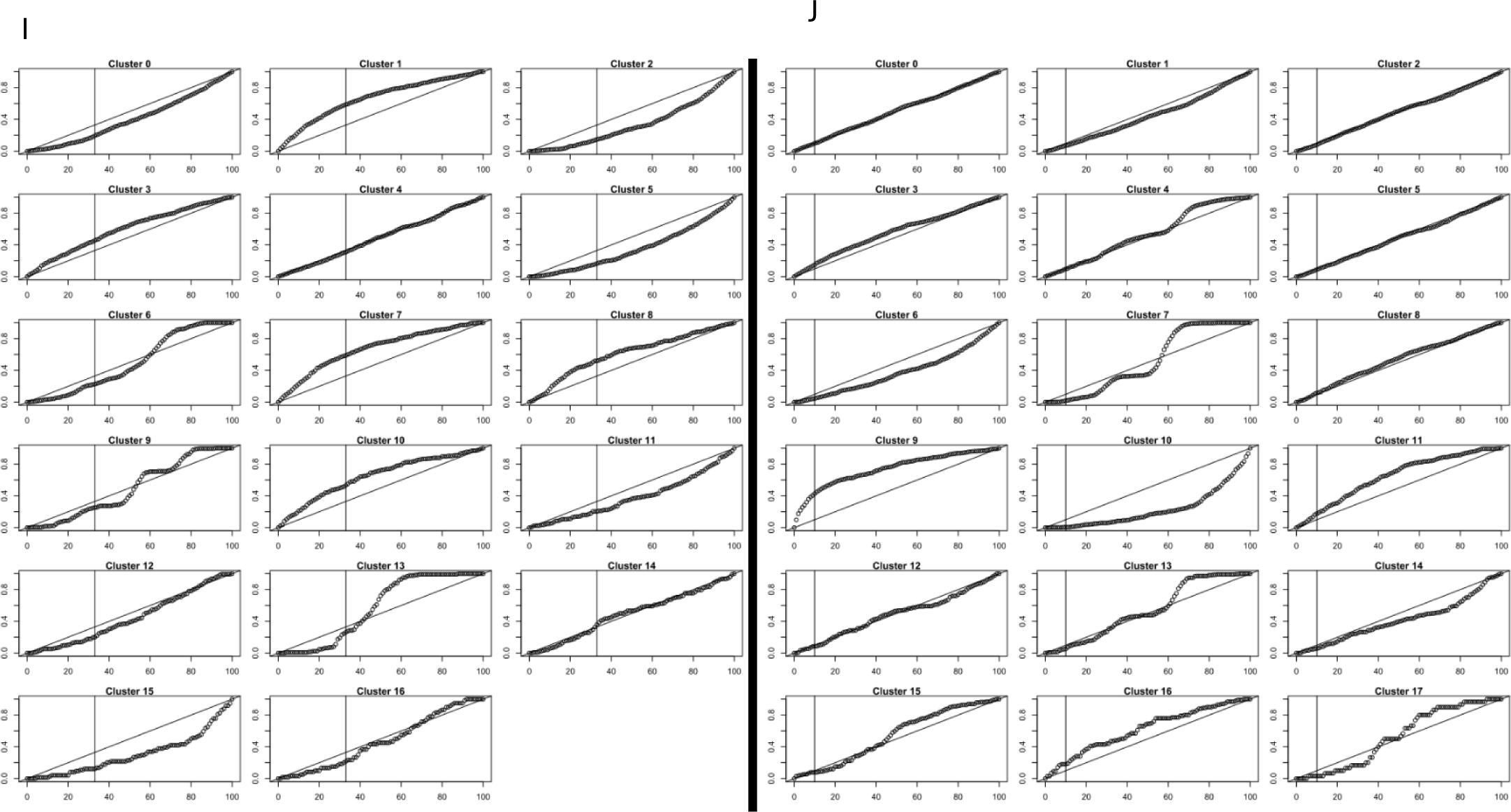
Intestinal module representation in each cluster. For each cluster in the male (left) or female (right) atlas the x-axis shows the fraction of the entire atlas being sampled in decreasing order of the score of the module being considered and the y-axis shows the fraction of the cluster that is represented in the sample. The vertical line shows the cutoff where the module scores equal 0 and the diagonal line shows linear growth (slope = 1). Clusters whose fractional representation increases faster than linear (above the diagonal line) in positive module scores (to the left of the vertical line) are clusters that contain high scoring cells for that module at a frequency higher than expected by chance alone (See Section ‘Annotating clusters: putative intestine’). The modules shown are: A) and B) unique genes from (Blazie et al., 2015), C) and D) genes from (Haenni et al., 2012), E) and F) enriched genes from (Kaletsky et al., 2018), G) and H) unique genes from (Kaletsky et al., 2018), and I) and J) genes from literature (McGhee, 2007) (See Section ‘Sources of marker gene modules’ for description of origins of intestinal modules).

Interestingly, yolk production is known to happen only in the hermaphrodite intestine of *C. elegans* and not in the male intestine (Perez & Lehner, 2019). The main protein component of the yolk, the vitellogenins are encoded by six genes *vit-1* to *vit-6* (Perez & Lehner, 2019). Three of these genes (*vit-3*, *vit-4*, and *vit-5*) have no orthologs in *H. bakeri*, while the remaining three have an ortholog with paralogs (HPOL_0001165701, HPOL_0001165801, and HPOL_0002023901). None of these *H. bakeri* genes are widely detected in either the male or female atlas. While yolk production would not be expected in the male intestine, the lack of transcripts for these proteins in the female atlas suggests that either few intestine cells were recovered in the female samples, and/or that *H. bakeri* do not make yolk, and/or that these orthologs have different functions to their *C. elegans* counterparts and the true yolk proteins remain unidentified in *H. bakeri*. Interestingly, the four vitellogenins in *A. suum* were found to be highly transcribed, even in male intestinal sections (Gao et al., 2017).

The LIGER cross-species analysis (Table S3) clusters cells of male cluster 1 and 3 with germline cells of *C. elegans*, while cells of male clusters 7 and 10 are clustered with somatic gonad and germline cells of *C. elegans*, and cells of male cluster 8 are clustered with somatic gonad, germline, and intestine cells of *C. elegans*. Cells from female cluster 3 clustered with germline cells of *C. elegans*, while cells of female cluster 9 clustered with germline, somatic gonad, egg-laying apparatus, and body wall muscle cells of *C. elegans*.

Taken together, these results suggest that male clusters 1, 3, 7, 8, and 10 are putatively intestine-related. The results are less clear for the female clusters. Of note, some of the female clusters that are represented by cells with high intestinal module scores are putatively oocyte/early embryo associated. It is unclear whether this reflects known connections (in *C. elegans*) between the intestine and the gonad (when yolk and other material is moved from intestinal cells to oocytes via receptor-mediated endocytosis), or whether newly forming intestinal tissue in early embryos begins to show common transcriptional signatures with adult intestine that quickly, or some other confounding factor. Female clusters not associated with oocyte/early embryo transcriptional patterns that may be potentially intestine associated include clusters 9, 3, and 15. Female cluster 15 also shows evidence of hypodermal-like transcription (in different cells within the cluster) and may be reflecting adult cells clustering separately from gamete and embryonic cells, which are quite common in the overall atlas. If this is the case, sub-clustering of cluster 15 may resolve different adult cell profiles, though there would still be few of them in the overall atlas.

#### Annotating clusters: putative neurons

Adult hermaphrodite *C. elegans* have 302 neurons representing 37% of the somatic cells by number (Hobert, 2010), yet being the smallest tissue by volume (Froehlich, Rajewsky & Ewald, 2021). Because of their characteristic long shape, they may not flow well through the microfluidic channels of the Chromium system, though 10X Genomics notes that adherent cells like neurons contract in solution which may allow their successful recovery (https://kb.10xgenomics.com/hc/en-us/articles/218170543-What-is-the-range-of-compatible-cell-sizes-). Therefore, to identify potential neuron cells we used the *H. bakeri* orthologs of the set of genes found to be unique to neurons (616 genes) from a *Punc-119::gfp* reporter strain (Kaletsky et al., 2018). The results suggest that male cluster 16 (≤ 18 cells) and female cluster 15 (≤ 22 cells) putatively include neurons. Both of these clusters are also associated with hypodermis expression profiles (and female cluster 15 has cells that score high in intestinal modules), albeit in different cells within the cluster. This may reflect similarities between the hypodermis and neurons, or may be a consequence of the small number of hypodermal cells and neurons that were recovered in each atlas not providing enough of an aggregate expression profile to accurately cluster cells of these two tissue types. Of note, cells from both of these clusters are clustered with neurons of *C. elegans* in the LIGER cross-species analysis (Table S3).

#### Annotating clusters: putative pharyngeal muscle

The pharynx in *C. elegans* consists of 95 cells of seven different types, including muscle, neurons, and epithelial cells (Kormish, Gaudet & McGhee, 2010). Despite these different cell types having common gene expression patterns with other similar cell types (ex. Pharygenal muscle and body muscle or pharyngeal neurons and tail neurons), there are also expression patterns common to the pharynx area, despite the cells being of different types (ex. The transcription factor PHA-4 is key for pharynx identity) (Kormish, Gaudet & McGhee, 2010). Assuming a similar phenomenon in *H. bakeri*, cells of the pharynx may or may not cluster with other cells of the same type or with other cells of the pharynx. Ideally each pharyngeal cell type would cluster on its own (as happened with adult *C. elegans* (Ghaddar et al., 2022)), but given that some of the cell types involved may or may not be represented well in the atlas (see putative neurons for example), pharyngeal cell identities may be hard to resolve. While *pha-4* does have an ortholog in *H. bakeri* (HPOL_0000795501), this transcript was undetected in the male atlas and detected in one cell in cluster 15 in the female atlas.

RNA-seq analysis from mixed-stage worms, including RNA bound to a polyA-binding protein that was expressed under the control of a pharyngeal muscle promoter (*myo-2p::PolyA-Pull*), defined a set of 312 genes to be uniquely expressed in the pharyngeal muscle, relative to intestine and body wall muscle (Blazie et al., 2015). Using orthologs of these genes as a module (Blazie_pharynx_uniq), the module scores for pharyngeal muscle are plotted on the male and female UMAPs (Figure 16A and B) and analyzed the same way as the intestinal modules (Figure 16C and D). High scoring cells are scattered throughout both male and female datasets, with slight enrichment in male clusters 4, 5, and 16 and female clusters 0, 4, and 15. LIGER cross-species analysis (Table S3) clusters cells of male cluster 4 with germline and somatic gonad cells of *C. elegans*, while cells of male cluster 5 cluster with germline cells of *C. elegans*. Cells of female cluster 0 cluster with germline cells of *C. elegans*, while cells of female cluster 4 cluster with germline and somatic gonad cells of *C. elegans*. Taken together, these results suggest that male clusters 4, 5, and 16 putatively include pharyngeal muscle. They also suggest that female clusters 0, 4, and 15 putatively include pharyngeal muscle. Both male cluster 16 and female cluster 15 have been associated with other cell types, which could indicate some spatial signals (similar to PHA-4 for pharynx) could be affecting clustering of the cells.

**Figure 16.**
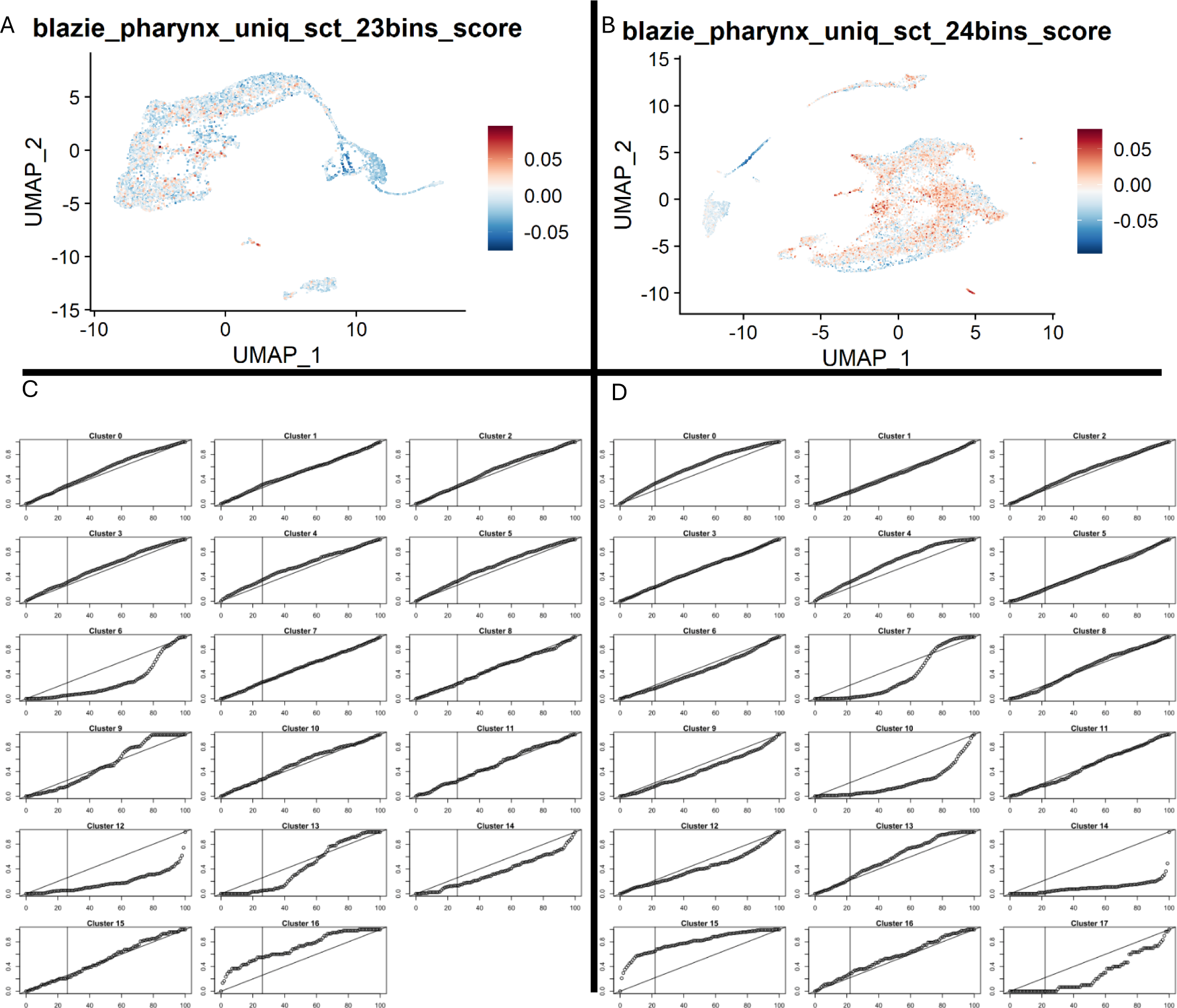
Pharyngeal muscle module scores. A) and B) cells are shown in either the male (A) or female (B) atlas, coloured according to their module score for the genes found to be unique to the pharynx in (Blazie et al., 2015). Module scores are relative and do not facilitate comparison to other modules or determination of a threshold score. C) and D) show the module representation in each cluster for the male (C) and female (D) atlases. The x-axis shows the fraction of the entire atlas being sampled in decreasing order of the scores for the module for genes unique to the pharynx (Blazie et al., 2015) and the y-axis shows the fraction of the cluster that is represented in the sample. The vertical line shows the cutoff where the module scores equal 0 and the diagonal line shows linear growth (slope = 1). Clusters whose fractional representation increases faster than linear (above the diagonal line) in positive module scores (to the left of the vertical line) are clusters that contain high scoring cells for that module at a frequency higher than expected by chance alone.

#### Annotating clusters: putative body muscle

In *C. elegans*, body wall muscles are the fourth largest tissue (Froehlich, Rajewsky & Ewald, 2021). Unlike vertebrate muscles, muscle cells in *C. elegans* are mononucleated, are completely post-mitotic, and have no satellite cells (stem cells) (Gieseler, Qadota & Benian, 2017). Assuming the same for *H. bakeri*, body wall muscles should be recovered well through the 10X Genomics Chromium for both adult males and females. A key marker gene in *C. elegans* muscle is the myosin gene *myo-3* (Gieseler, Qadota & Benian, 2017). The ortholog of this gene in *H. bakeri* (HPOL_0001848901) is detected in six cells in the male atlas (clusters 5, 11, and 16) and 99 cells of the female atlas (all clusters except 6, 7, 10, 14, and 17) (Figure S3). The extreme underrepresentation of *myo-3*-expressing cells in either dataset is unexpected and suggests muscle cells were not recovered well.

Orthologs of two sets of potential body-muscle-specific genes in *C. elegans* were used as modules to try to better resolve any recovered body muscle cells. The first set (Blazie_bodymuscle_uniq) was 329 genes uniquely expressed in body muscle, relative to pharyngeal muscle and intestine, obtained from sequencing from mixed-stage worms RNA bound to a polyA-binding protein that was expressed under the control of a body muscle promoter (*myo3*) (Blazie et al., 2015). The second set was the set of genes found to be unique to body muscle in adults from cell sorting of a *Pmyo-3::mCherry* reporter strain (Kaletsky et al., 2018). The module scores for these two modules are plotted on the male and female UMAPs (Figure 17) and analyzed the same way as the intestinal modules (Figure 18). LIGER cross-species analysis clusters cells of male clusters 0 and 2 with germline cells of *C. elegans* (Table S3), while cells of male cluster 11 cluster with germline and hypodermis cells of *C. elegans*. Interestingly, LIGER clusters cells of female cluster 12 with either germline, body wall muscle, or egg-laying apparatus cells of *C. elegans*, with the most represented cell type (when only including the female *H. bakeri* sample) being uterine muscle. While this cluster may be associated with uterine muscle, we had no marker genes to use as additional evidence. Taken together, these results suggest that male clusters 0, 2, 4, 5, 11, and 16 putatively include body wall muscle. They also suggest that female clusters 0, 4, and 15 putatively contain body wall muscle. Based on the clusters putatively associated with pharyngeal muscle, there may be some clustering of muscle cells together based on common expression profiles, while there also seems to be some clustering of different cell types together (ex. Female cluster 15) based on other, unknown, signals.

**Figure 17.**
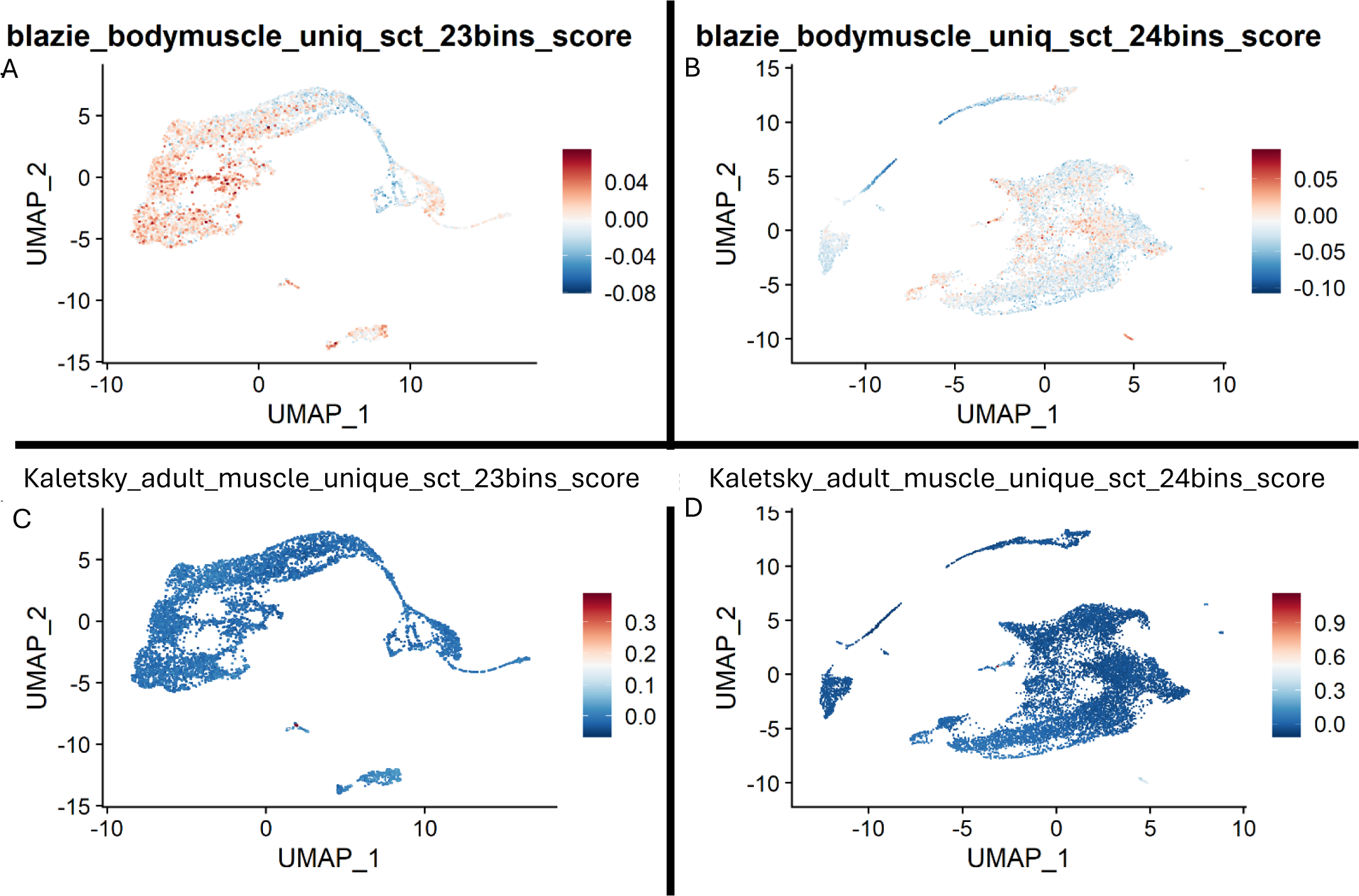
Body muscle module scores to identify potential muscle cells. Cells are shown in either the male (left) or female (right) atlas, coloured according to their score in the modules: A) and B) genes found to be unique to body muscle in (Blazie et al., 2015), or C) and D) genes found to be unique to muscle in (Kaletsky et al., 2018). Module scores are relative and do not facilitate comparison to other modules or determination of a threshold score.

**Figure 18.**
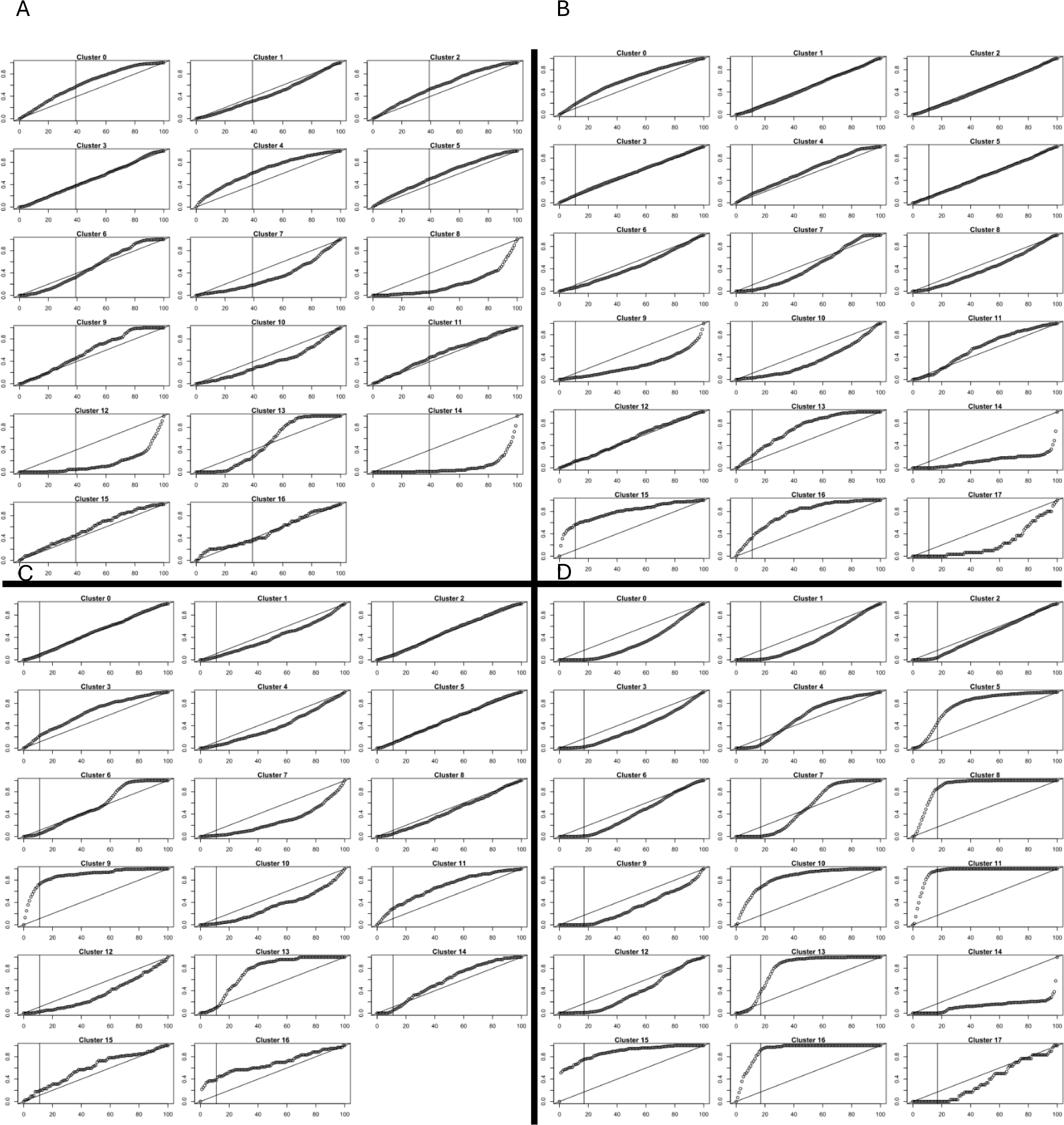
Body muscle module representation in each cluster. For each cluster in the male (left) or female (right) atlas the x-axis shows the fraction of the entire atlas being sampled in decreasing order of the score of the module being considered and the y-axis shows the fraction of the cluster that is represented in the sample. The vertical line shows the cutoff where the module scores equal 0 and the diagonal line shows linear growth (slope = 1). Clusters whose fractional representation increases faster than linear (above the diagonal line) in positive module scores (to the left of the vertical line) are clusters that contain high scoring cells for that module at a frequency higher than expected by chance alone. The modules shown are: A) and B) genes found to be unique to body muscle in (Blazie et al., 2015) and C) and D) gene found to be unique to muscle in (Kaletsky et al., 2018).

#### Marker genes for cluster ID verification and best practices for sample handling

The identities of the cells making up the clusters need to be verified empirically. This is commonly done in other scRNA-seq studies using hybridization-based methods, such as fluorescent in situ hybridization (FISH) as used in (Swapna et al., 2018), or whole-mount in situ hybridization (WISH) as used in (Wendt et al., 2020). These techniques use probes to target specific transcripts, which are selected from the marker genes identified in the clustering analyses. The method used by Seurat for finding marker genes selects features that are up-regulated in a cluster relative to the rest of the atlas. Without any requirement that these markers be uniquely up-regulated in the cluster being considered, the result is that any given feature can be a marker for more than one cluster. When two or more clusters that share a marker gene are made up of cells of the same type, this isn’t a problem with respect to using the markers to identify the cell type of the cluster(s). However, to best identify the clusters, marker genes that uniquely identify each cluster are preferred. Therefore, from the full list of marker genes predicted for every cluster in each atlas (Tables S1 and S2), we retrieved the genes that were unique for a single cluster and ordered them by expression level as candidate markers for follow up cluster identification experiments (Tables S36 and S37).

Based on the putative cluster annotations (see above sections), there is evidence that many of the abundant cell types in the worms were recovered at some level, even if not resolved into discrete clusters or recovered at the level expected based on cell-type abundance in *C. elegans*. A comprehensive scRNA-seq atlas of adult *C. elegans* contains ∼150,000 cells (Ghaddar et al., 2022). We randomly subsampled this atlas to contain 11,000 cells (roughly the size of the *H. bakeri* female atlas) and 6,000 cells (roughly the size of the *H. bakeri* male atlas) and reanalyzed it using monocle3 (the same software used in the original analysis) to see if the originally identified cell types could be recovered as discrete clusters with significantly fewer cells included in the analysis (Tables 4, S38, and S39). At 6,000/150,000 cells, missing cell groups include rectum, head mesoderm, GLR, excretory, and embryonic cells (even though these cell types are included in the analysis), indicating that cells of different types cluster together when the number of cells in the analysis is too low. In the *H. bakeri* datasets there are clusters that appear to be multiple cell types clustering together, indicating that the number of cells recovered is insufficient. It is also likely that certain cell types were excluded by the methods used to dissociate the samples or sort the cells before running the Chromium because in the LIGER cross-species analysis there were several cell type groups from the Ghaddar groups that rarely, if ever, clustered with the *H. bakeri* cells recovered (ex. Coelomocytes, excretory, rectum, seam) (Table S3).

**Table 4.** Cell type groups from (Ghaddar et al., 2022) that would be recovered as discrete clusters at different sizes of their atlas.

| Cell type group | Full atlas (154,251 cells) | # clusters at 11,000 cells | # clusters at 6,000 cells |
| --- | --- | --- | --- |
| Atypical cells | 1 | 0 | 0 |
| Body wall muscle | 4 | 1 | 1 |
| Coelomocytes | 1 | 1 | 1 |
| Egg-laying apparatus | 8 | 2 | 2 |
| Embryonic | 1 | 0 | 0 |
| Excretory | 3* | 2 | 0 |
| Germline | 17 | 2 | 2 |
| GLR cells | 1 | 1 | 0 |
| Head mesodermal cells | 1 | 0 | 0 |
| Hypodermis | 4 | 1 | 1 |
| Intestine | 5 | 1 | 2 |
| Neurons | 85 | 12 | 6 |
| Pharynx | 9 | 4 | 1 |
| Rectum | 2 | 0 | 0 |
| Seam | 2* | 1 | 1 |
| Somatic gonad | 12 | 4 | 3 |
| Support cells | 9 | 1 | 1 |
| Unassigned | 6 | 0 | 0 |
| Total Clusters | 170 | 33 | 22 |

\*Note that one cluster is evenly split between hypodermis and seam cells

### Limitations of orthology-based methodology and biological differences between the worms

Since there are no verified cell type markers in *H. bakeri*, using predicted orthologs of markers from *C. elegans*, or other model organisms, is the only way to find putative cluster identities informatically. This is a critical step for assessing whether the parameters used in the analysis were properly tuned to yield the best possible atlas, and therefore the best possible clusters and cluster markers. These markers in turn are needed to verify the identities of the clusters empirically. However, since hybridization data to verify the cluster identities is not available, all downstream analyses, from this point in the document on, have had to rely on the putative informatic identities. Using orthologs of cell markers from other organisms in this manner is inherently assuming that the ortholog produces the same gene product, with the same function, and the same expression patterns, including temporal expression and tissue specificity. While these assumptions may be true for some ortholog pairs, there are examples even from within the present analysis where they are not true (ex. *spe-4*, which is a sperm-specific presenilin in *C. elegans*, but whose expression is not strictly restricted to the sperm in the *H. bakeri* atlases).

Moreover, ortholog prediction itself is an imperfect process (Natsidis et al., 2021), adding additional noise to the putative cluster identifications. Of note, using orthologs to *C. elegans* markers to identify scRNA-seq clusters was also attempted with *Brugia malayi*, with similar limited success (61.5% metaof total cells remain unannotated) (Henthorn et al., 2023).

Using the orthology information, from WormBase ParaSite, between *C. elegans* and *H. bakeri* does, however, highlight some key similarities and differences between these two closely related worms. Assuming all the clusters are correctly identified informatically, it appears that spermatogenesis occurs similarly in the two worms, with spermiogenesis occurring within the female after mating. While several key genes expressed in *C. elegans* sperm had no ortholog in *H. bakeri* (ex. the paternal effect lethal gene *spe-11*), there were enough orthologs of sperm-marker genes to implicate clusters in this analysis as sperm-related, putatively right down to the level of the stage of differentiation. Likewise, it appears that the eggshell is superficially similar between the two worms. Not only did genes related to the layers of the *C. elegans* eggshell help to putatively identify oocyte/early embryo clusters, but the much better digestion of the eggshell in the pronase samples than the liberase samples (and the chitinase activity of pronase but not liberase) strongly argues for a key chitin layer in the *H. bakeri* eggshell, as in *C. elegans*. The size of the clusters putatively associated with hypodermis suggests that either the hypodermis in *H. bakeri* is almost unrecognizable to the hypodermis of *C. elegans*, or that, like *C. elegans*, most of the hypodermis forms a large syncytium that was not recovered well after sample processing. The latter is likely the case since part of the identification of hypodermis was based on cuticle collagens of *H. bakeri* and not on direct orthology to *C. elegans* genes. Similarity of the hypodermis between the two worms, and the expression of orthologs of other cuticle components (see hypodermis section) suggests that the structure and function of the cuticle of the two worms are at least coarsely similar.

On the other hand, the computed cell cycle scores and putative cluster identifications in the *H. bakeri* atlases suggest that cells other than the germline are still actively cycling (Figure 2C and D). In *C. elegans*, the adult somatic cells do not divide and are not actively cycling (Hubbard & Schedl, 2019). In addition to cells categorized as S phase or G2M phase by Seurat’s CellCycleScoring method being well represented in putatively somatic clusters (ex. male cluster 11 or female cluster 3), we analyzed the assigned cell cycle phase of individual cells with the highest scores in certain modules (Table 5). In the male atlas, where there was greater recovery of putative adult somatic tissues, both the intestine and muscle tissues appear to be cycling and dividing, while hypodermis and neurons do not. Mitotic divisions have been observed in adult *A. suum* intestine at a rate of 0.01-0.1 divisions per 1000 cells and were found to account for 86% of the adult growth of the worm (Anisimov & Tokmakova, 1974; Anisimov & Usheva, 1974). Actively cycling and dividing somatic cells in *H. bakeri* would represent a major difference between the two worms, if confirmed to be the case.

**Table 5.** Assigned cell cycle phase in different adult *H. bakeri* somatic tissues.

| Tissue | Module | Score cutoff | Quantile | # cells | # cells in G1 phase | # cells in G2M phase | # cells in S phase |
| --- | --- | --- | --- | --- | --- | --- | --- |
| Male intestine | literature | > 0.05 | Top 4% | 220 | 127 | 74 | 19 |
| Male body muscle | Blazie bodymuscle uniq | > 0.05 | Top 1% | 36 | 8 | 14 | 14 |
| Male pharyngeal muscle | Blazie pharynx uniq | > 0.05 | Top 0.5% | 24 | 5 | 12 | 7 |
| Male hypodermis | Kaletsky adult hypodermis uniq | > 0.1 | Top 0.5% | 20 | 20 | 0 | 0 |
| Male neurons | Kaletsky adult neuron uniq | > 0.1 | Top 0.5% | 14 | 12 | 2 | 0 |

### Adult male average intestinal expression profiles

The average expression values for all clusters for both the male and female atlases can be found in Table S6 – S9. Since the male atlas has more clusters that are putatively adult tissues, the adult profiles are described from the male atlas only. The adult male intestine is putatively contained in male clusters 1, 3, 7, 8, and 10. These clusters are highly transcriptionally active (Table 6), GO enrichment results of all transcripts detected in these clusters and of the highly expressed transcripts in these clusters can be found in Tables S12–S21. Broad activities within these clusters include protein synthesis, maintenance of amino acid and nucleic acid pools, transport, energy generation, and catabolism. Highly expressed transcripts reflect protein synthesis, energy generation, and biosynthetic processes (Tables S17–S21). By comparing each of these clusters to all the remaining clusters in the atlas, 129 transcript features were found to be significantly up-regulated in all intestinal clusters and 103 were found to be significantly down-regulated in all intestinal clusters (Tables S10 and S11). Among the consistently up-regulated genes are genes involved with calcium storage and regulation, including the ortholog of calreticulin (*crt-1* in *C. elegans*), a gene expressed in the intestine and important for the defecation cycle in *C. elegans* (McGhee, 2007), a calcium-binding EF-hand domain protein, a store-operated calcium entry-associated regulatory factor, and a bax-inhibitor 1-related protein. Additionally, there are genes involved in vesicular trafficking, amino acid metabolism, and fatty acid metabolism. There are also two potential transcription factors (HPOL_0000751101 and HPOL_0001055401), which may be particularly important for regulating intestinal functions. Notably, HPOL_0001055401 has no ortholog in *C. elegans* (though does have orthologs in other nematodes), while HPOL_0000751101 is categorized as an ortholog *lin-1* in *C. elegans*, which has been demonstrated to be involved in vulval formation (Beitel et al., 1995). Finally, there are genes potentially involved in protein secretion, including SecY/SEC61-alpha family, TRAM1-like, Protein translocase complex, SecE/Sec61-gamma subunit, translocon-associated, and signal peptidase-like proteins. Among the consistently down-regulated genes are genes specific to the function of other tissues (ex. major sperm proteins, macoilins that are involved in neuronal functions, etc.), as well as a putative sugar/inositol transporter (HPOL_0000113301), and a major intrinsic protein (HPOL_0001535701) that may function in water transport. There is also a transthyretin-like protein (HPOL_0001855101), which are nematode secreted proteins, suggesting this protein is produced and secreted elsewhere.

**Table 6.** Stats of average male cluster profiles.

| Cluster | # transcript features detected (> 0) | # highly expressed transcripts (> 1.7415386266, top 10%) |
| --- | --- | --- |
| 0 | 11159 | 874 |
| 1 | 11256 | 1201 |
| 2 | 10984 | 867 |
| 3 | 11090 | 1012 |
| 4 | 11000 | 1343 |
| 5 | 11568 | 989 |
| 6 | 7098 | 788 |
| 7 | 10948 | 1159 |
| 8 | 10452 | 1094 |
| 9 | 5891 | 998 |
| 10 | 9439 | 1179 |
| 11 | 12649 | 1028 |
| 12 | 9789 | 1159 |
| 13 | 6090 | 205 |
| 14 | 9466 | 1046 |
| 15 | 8418 | 935 |
| 16 | 9393 | 792 |

To further examine activities that may be localized to different parts of the intestine, and to explore the differences between the five putative intestinal clusters, we compared the gene expression between the intestinal clusters (Tables S22–S31). Clusters 3 and 8 appear to be the most different from each other based on the number of significantly differentially expressed genes in each pairwise comparison (Tables S22–S31). In all pairwise comparisons, GO terms of genes that are significantly up-regulated in cluster 3 reflect translation and biosynthetic pathways, while GO terms of genes that are significantly up-regulated in cluster 8 (particularly relative to cluster 3) reflect catabolism and localization. The calcium-related genes mentioned above show the highest expression in cluster 8 with decreasing expression in clusters 7 and 10, then cluster 1, and lowest expression in cluster 3. Given that genes associated with the defecation cycle in *C. elegans* are more highly expressed in the posterior intestine, where the cyclic calcium fluctuations initiate (McGhee, 2007), this tentatively suggests that cluster 8 represents posterior intestinal cells, with clusters 7 and 10, then cluster 1 being more anterior, and cluster 3 being the most anterior intestinal cells. This would therefore suggest that the anterior intestine is more focused on protein synthesis (potentially of digestive enzymes), while the posterior intestine performs more of the catabolism of the acquired nutrients and localizes the macromolecules accordingly. Likewise, bulk RNA-seq analysis of anterior, middle, and posterior sections of the *A. suum* intestine found specialization of function along the anterior-posterior axis of the intestine and suggested a larger role for the middle intestine in performing biosynthetic functions (Gao et al., 2017).

I further examined the expression patterns of the cytochrome P450 genes in *H. bakeri*. Members of this large gene family are involved in general metabolism and implicated in drug metabolism in other parasitic nematodes (Laing et al., 2013). *H. bakeri* has 33 genes that are annotated with the Interpro domain for the cytochrome P450 superfamily (IPR036396). Of these, only one is significantly differently expressed in the intestine relative to the non-intestinal clusters: HPOL_0000554501 is significantly down-regulated in all intestinal clusters relative to non-intestinal clusters. This gene is an ortholog of the *C. elegans* gene *cyp-37B1*, which has been implicated in response to ivermectin exposure and is expressed in the intestine in *C. elegans* (Laing et al., 2012). Part of the expression pattern observed here in *H. bakeri* is being driven from the high expression of this gene in the sperm-related clusters (Figure 19), for which this gene is a marker.

**Figure 19.**
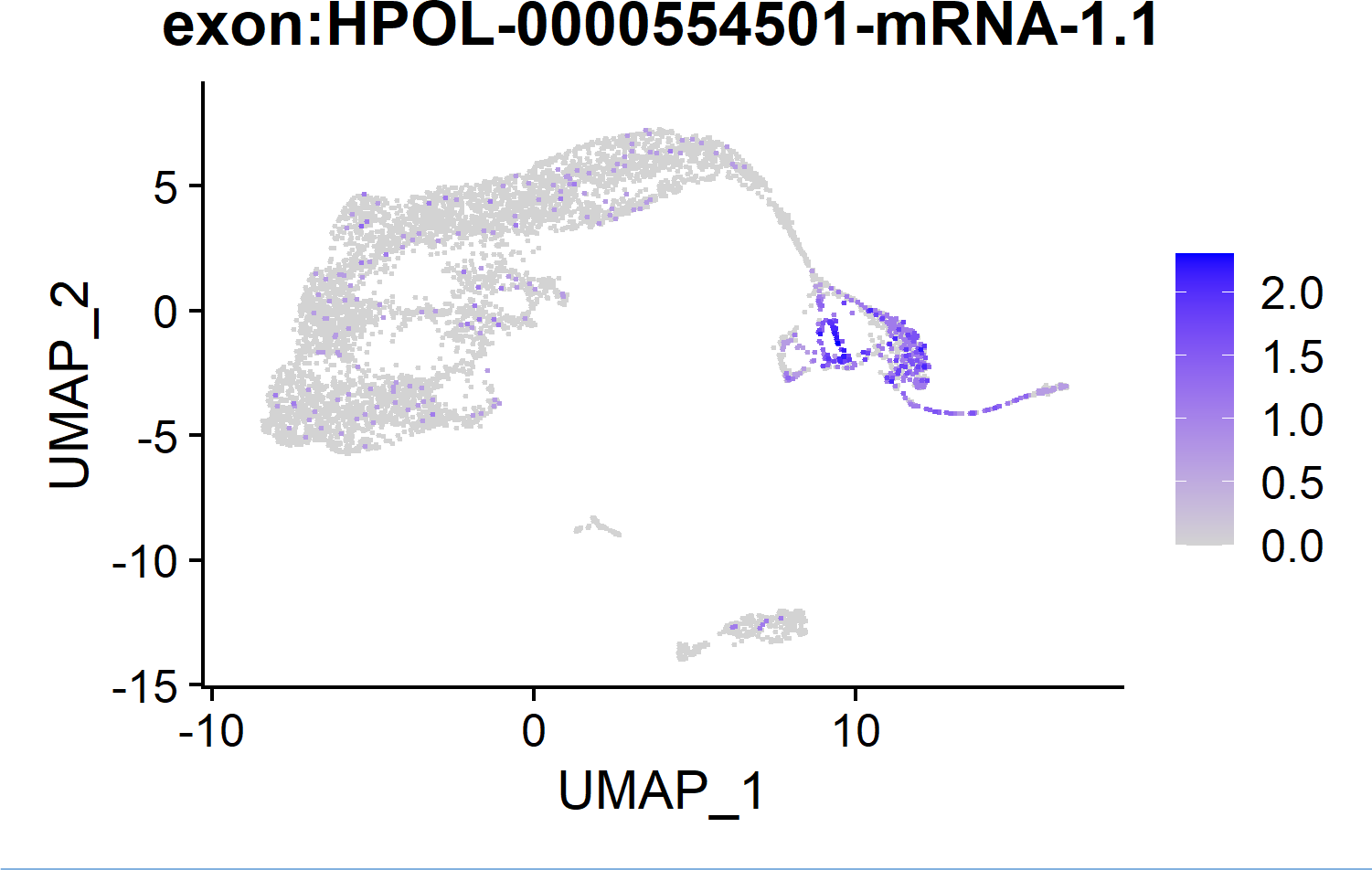
FeaturePlot of expression of the *H. bakeri* ortholog of *cyp-37B1* in *C. elegans*. Cells in the male atlas are coloured according to their expression level (SCT-normalized UMI count for that transcript for that cell).

### Early embryogenesis in *H. bakeri* vs *C. elegans*

When a spermatozoon fertilizes an oocyte, the contents of the two cells join, resulting in a one-cell embryo whose transcripts are entirely of parental origin. To define which transcripts are contributed by each parent to the resulting embryo, and to try to use the sperm information to further resolve the oocytes from the early embryos, we examined the transcript features shared between putative sperm clusters (male cluster 13, female cluster 14, and female cluster 7) and putative oocyte or newly fertilized embryo clusters (female clusters 13, 16, 11, and 8). Ideally (if we could know the true sperm profile, the true oocyte profile and the true one-cell embryo profile), we would expect to see that the one-cell embryo profile would contain all of the oocyte profile and more of the sperm profile than what the oocyte profile has. We would also expect the lowest overlap between the sperm and oocyte profiles, while the oocyte profile would have much of the one-cell embryo profile and decreasing similarity with the profiles of embryos further along in their development (especially after the PZT). Whether based on features detected or features above a certain expression threshold, the proportions of features shared between the sperm and oocyte/embryo clusters (Figures 20, S4, S5 and S6, Table S32) suggest that female cluster 16 represents unfertilized oocytes (lowest overlap between sperm profiles and c16, orange bars of bottom 3 panels of Figure 20), female cluster 13 contains newly fertilized one-cell embryos (c16 contains proportionally more features of c13 than c11 or c8, green bars in first panel of Figure 20 and c13 contains greater proportion of sperm features than c16, green bars in first two panels), and female clusters 11 and 8 contain embryos that have begun the PZT. Consequently, maternal contributions to the embryo can be defined as features that are common to female cluster 16 and 13 and paternal contributions as features common to female cluster 13 and a sperm profile (female cluster 7, or combined female clusters 7 and 14 and male cluster 13). When basing parental contributions on features detected (i.e., expression greater than 0 in the cluster), transcripts from 5267 genes are contributed to the embryo from the mother (MCO.ALL) and transcripts from 4993 genes are contributed to the embryo from the father (PCO.ALL) (Table S33). Interestingly, 4222 of these genes are common to both the maternal and paternal contributions (PAR.SHARED), leaving 771 potentially uniquely paternal contributions (PCO.U) and 1045 potentially uniquely maternal contributions (MCO.U). Of note, it has been found in *C. elegans* that not all transcripts present in the sperm end up in the embryo, suggesting a selection of mRNAs that are transferred during fertilization (Stoeckius, Grün & Rajewsky, 2014), which, if also true in *H. bakeri*, would suggest that most of the 4222 PAR.SHARED features, though present in the sperm, are being contributed to the embryo by the oocyte.

**Figure 20.**
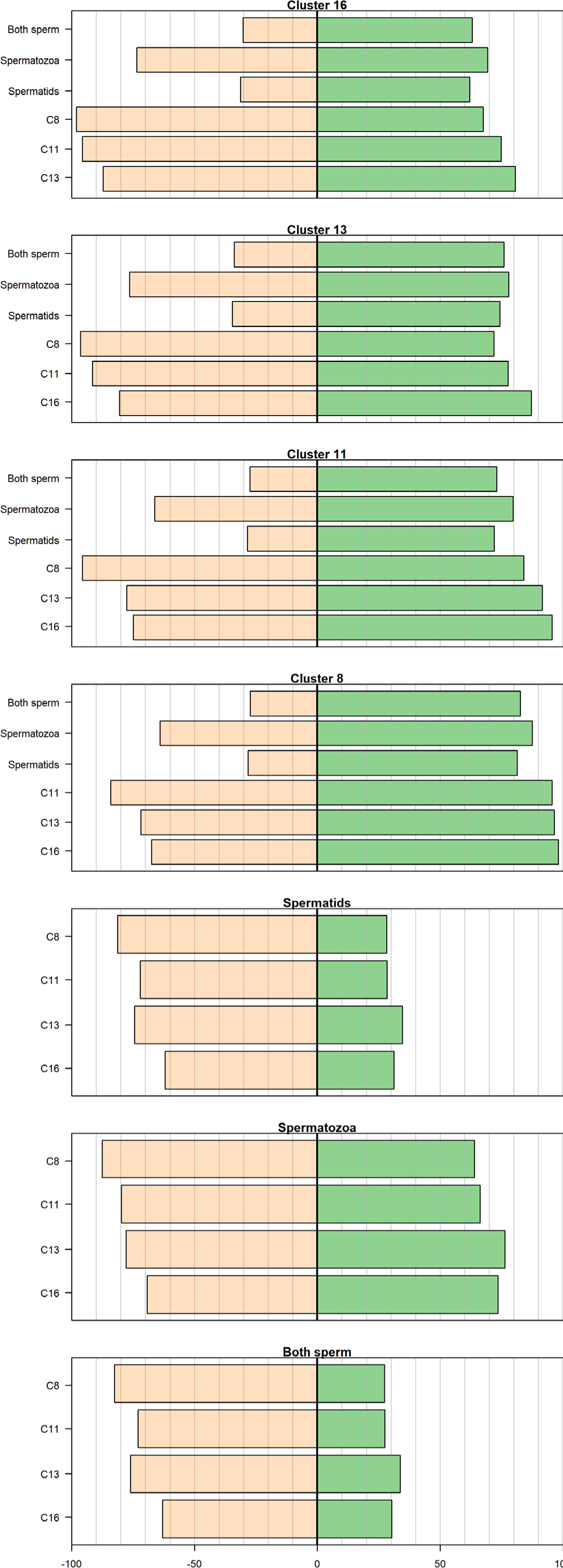
Percentage of transcript features shared among putative sperm and oocyte/embryo clusters. See Section ‘Early embryogenesis in *H. bakeri* vs *C. elegans*’ for details. In each panel green bars denote the percentage of features in the cluster labelled on the left that are found in the cluster in the title, while orange bars show the percentage of features in the cluster in the title that are found in the cluster labelled on the left. Grey lines line up to the x-axis at the bottom and mark every 10%. The black middle line is 0. Features were considered found in a cluster if they were detected as expressed at all in the cluster (expression > 0).

Additionally, RNAPII is silent in *C. elegans* early embryos (Baugh et al., 2003; Stoeckius, Grün & Rajewsky, 2014) and the sperm have been found to contribute ∼10% of the RNA to the embryo in *C. elegans* (Stoeckius, Grün & Rajewsky, 2014). Assuming the same in *H. bakeri*, features that are upregulated in the 1-cell embryo relative to the oocyte are good candidates for being paternal contributions (PCE). DGE between female cluster 13 and 16 results in 617 features that are significantly (p_adj_ < 0.05) up-regulated in the one-cell embryo (cluster 13) relative to the oocytes (cluster 16) (Table S33). Of these 617 PCE features, 10 are also among the 771 PCO.U features defined above. These 10 genes and their annotations are listed in Table 7. The remaining 607 PCE features are all found in the 4222 PAR.SHARED features that are common to both maternal and paternal contributions.

**Table 7.**
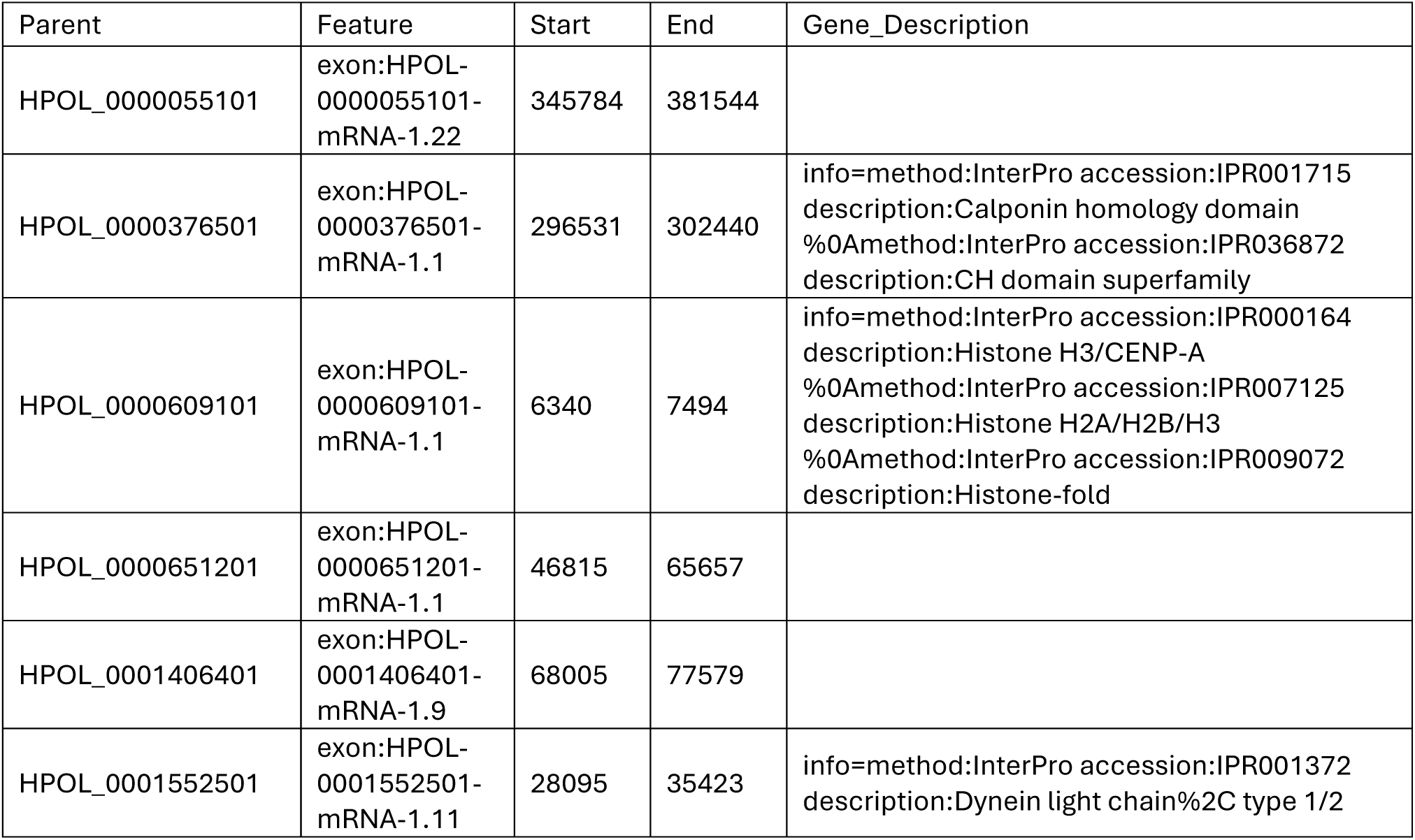
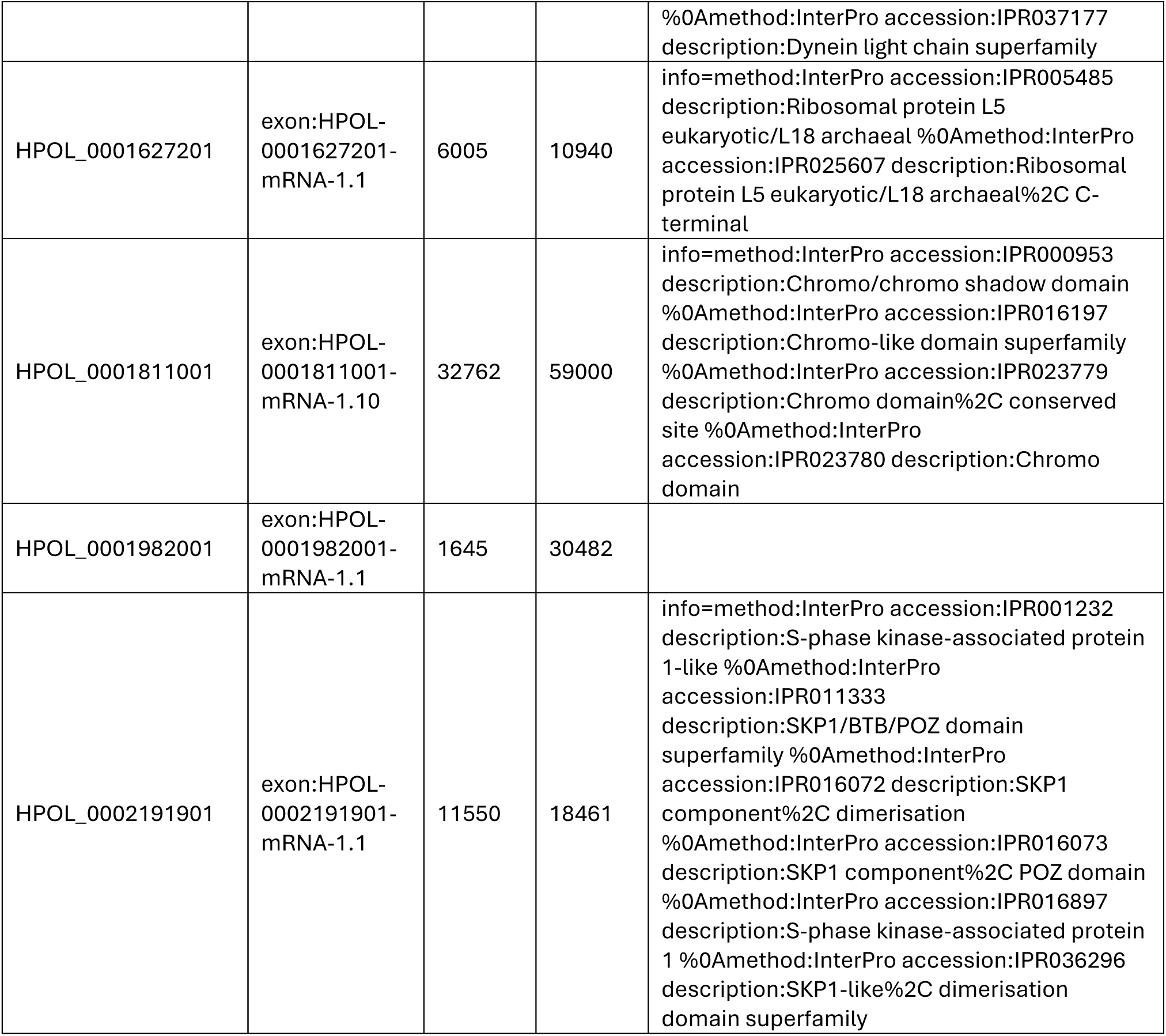
Paternal contributions to the embryo, defined by both overlapping features and differential expression.

In comparison, in *C. elegans*, 164 genes have been identified as paternal contributions to the embryo, though the authors note that their method is under-estimating the true number, with 60% of potential paternal contributions not discoverable (Stoeckius, Grün & Rajewsky, 2014). Moreover, it was noted that paternal contribution transcripts are not necessarily highly expressed and are largely uncharacterized with no functional information, but for those for which there was functional information there was an enrichment of genes involved in embryonic lethal and maternal sterile phenotypes (Stoeckius, Grün & Rajewsky, 2014). Here in *H. bakeri*, the 771 PCO.U potentially paternal contributions feature 32 enriched GO terms involving protein kinase and phosphatase activity (Table S34), while the 617 PCE potentially paternal contributions feature 124 enriched GO terms involving RNA processing, metabolic, and oxidative activities (Table S35). Of the 771 PCO.U potentially paternal contributions, 310 have no annotation information, while of the 617 PCE potentially paternal contributions, 66 have no annotation information, highlighting the understudied nature of paternal contributions to early embryonic development. Of the 164 paternally enriched genes in *C. elegans*, 94 have an ortholog in *H. bakeri*. Of these 94 genes, three are found in the 771 PCO.U potentially paternal contributions and 5 are found in the 617 PCE potentially paternal contributions, with none found among the 10 genes common to the two sets. Taken together these results point to many (unknown) differences in early development between the two worms.

### Conclusions

Though this first attempt at scRNA-seq in *H. bakeri* did not yield a complete atlas, due to sample processing upstream of the Chromium, it does still afford an opportunity to compare gene expression in specific cell types between *C. elegans* and *H. bakeri*. The putative identities of the cells robustly recovered here include the gametes of both sexes, embryos of various stages, and adult male intestine, while hypodermal, muscle, neural, and pharyngeal cells are under-represented and/or co-clustering. Putatively identifying cell types in *H. bakeri* using orthologs of genes in *C. elegans* suggests that the two worms have a coarsely similar hypodermis, cuticle, and eggshell, as well as spermatogenesis process. It also suggests that, unlike in *C. elegans*, intestinal and muscle cells in *H. bakeri* may still be actively cycling and dividing. Within the intestine, there appears to be a spatial segregation of intestinal functions along the anterior-posterior axis, with the anterior focused more on protein synthesis and the posterior more on catabolism and macromolecule localization. Finally, early embryogenesis and development appears to be very different between the two worms, with only eight of 94 confirmed *C. elegans* paternal contributions (with an ortholog) also potentially being paternal contributions in *H. bakeri*.

## Funding

Doctoral scholarships from the Killam Trust and Natural Sciences and Engineering Research Council of Canada (NSERC) to SMJP.

Grants from NSERC, Results Driven Agricultural Research (RDAR, Alberta), and Alberta Innovates Technology Futures (AITF) to JDW.

Grant from NSERC to CAMF.

The Calgary Firefighters Burn Treatment Society Chair to JB.

## Acknowledgements

We thank Drs. Stephen Doyle, John Gilleard, David Hansen, Dongyan Niu, and Tarah Lynch for their help and their comments on earlier drafts of this manuscript. We acknowledge the high-performance computing resources made available by the Faculty of Veterinary Medicine and Research Computing at the University of Calgary.

## Supplementary Material

Supplementary tables can be found in a single excel file with every table in its own sheet.

### Supplementary Figures

**Figure S1.**
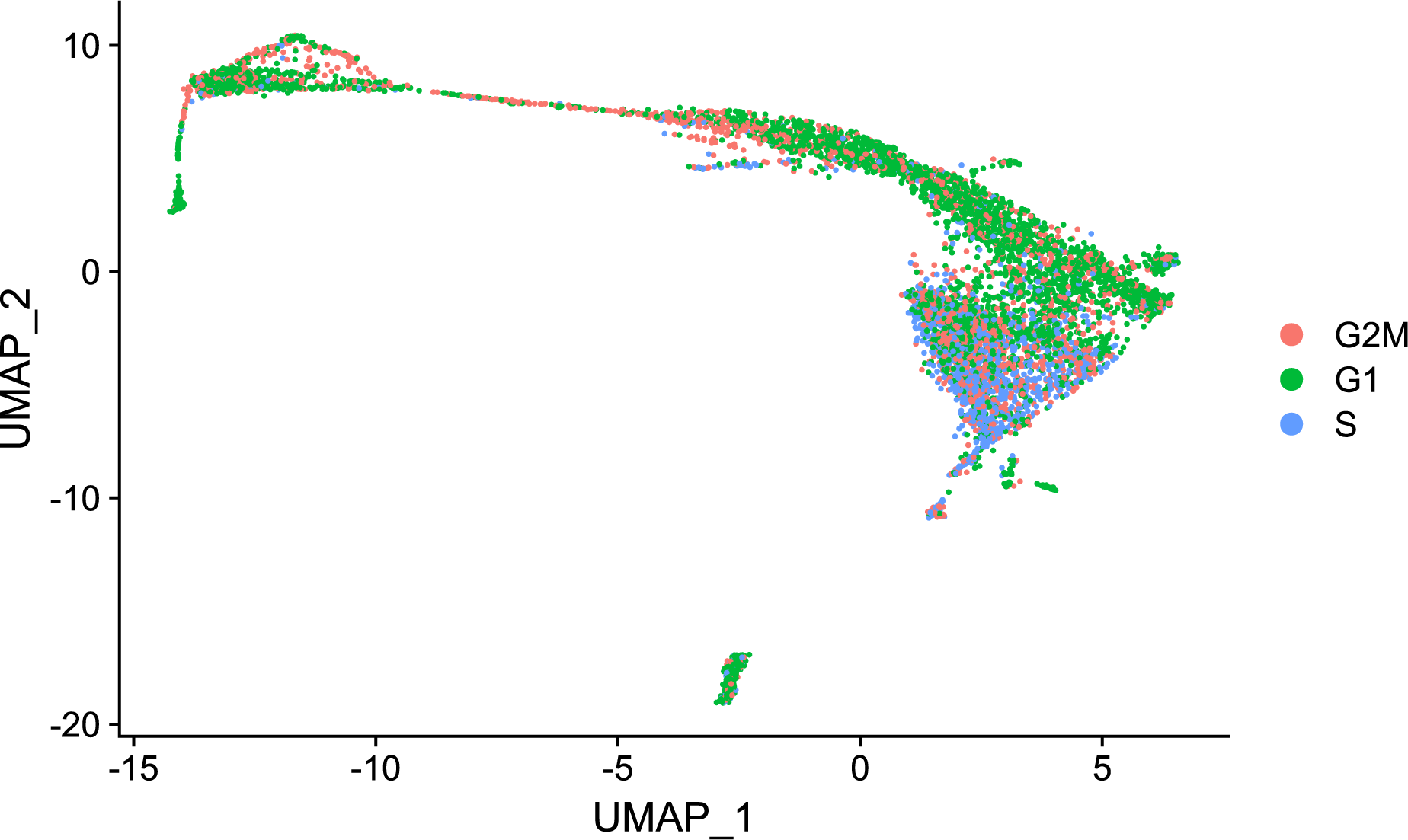
UMAP of *H. bakeri* single-cell male atlas with no cell cycle regression during normalization. Cells are coloured according to the cell cycle phase they were assigned by Seurat.

**Figure S2.**
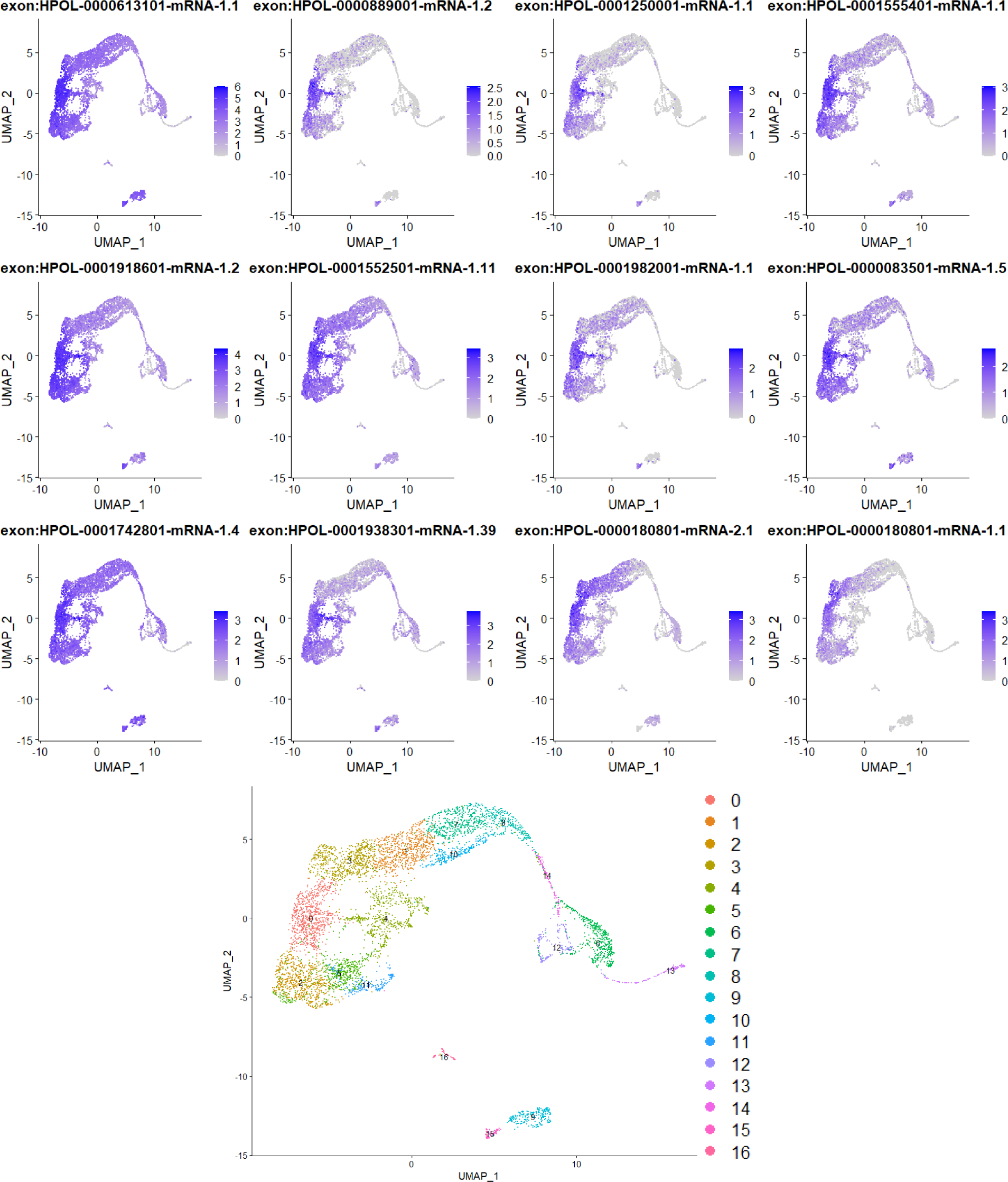
Example expression patterns for the top twelve cluster markers of male atlas cluster 0. Expression levels of the top twelve cluster markers for male cluster 0 (when ranked by increasing adjusted p-value) were used to colour the cells on a UMAP of the male atlas. The UMAP of the male atlas is also shown with cells coloured by cluster assignment.

**Figure S3.**
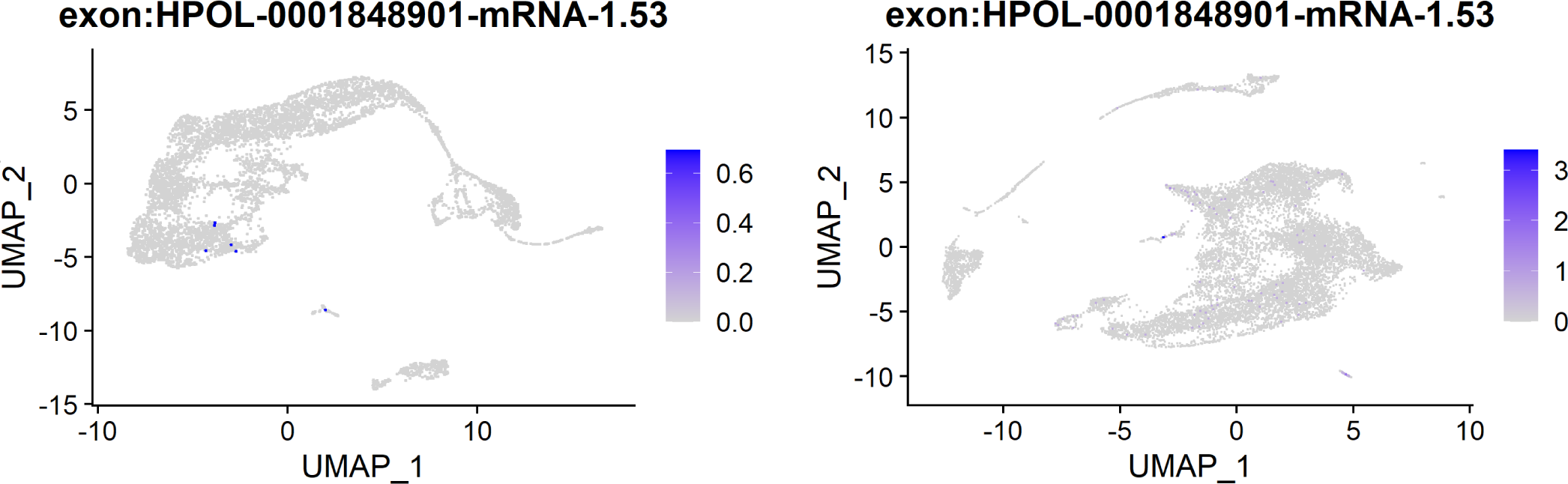
FeaturePlots of the expression of the ortholog of *myo-3* in the male (left) and female (right) atlases. Cells are coloured according to the expression level of the *H. bakeri* ortholog of *myo-3*.

**Figure S4.**
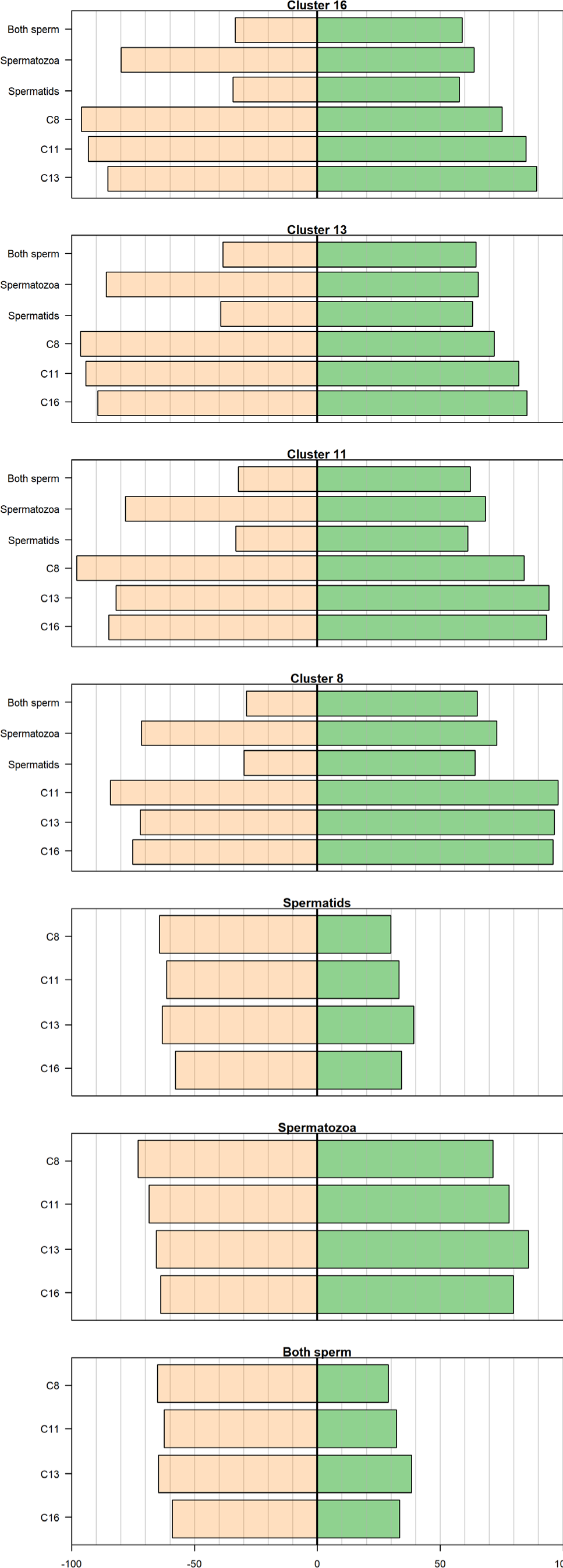
Proportions of features shared among putative sperm and oocyte/embryo clusters. In each panel green bars denote the percentage of features in the cluster labelled on the left that are found in the cluster in the title, while orange bars show the percentage of features in the cluster in the title that are found in the cluster labelled on the left. Features were considered found in a cluster if their average cluster expression value was > 0.01, which corresponds to the top 82% of feature expression values.

**Figure S5.**
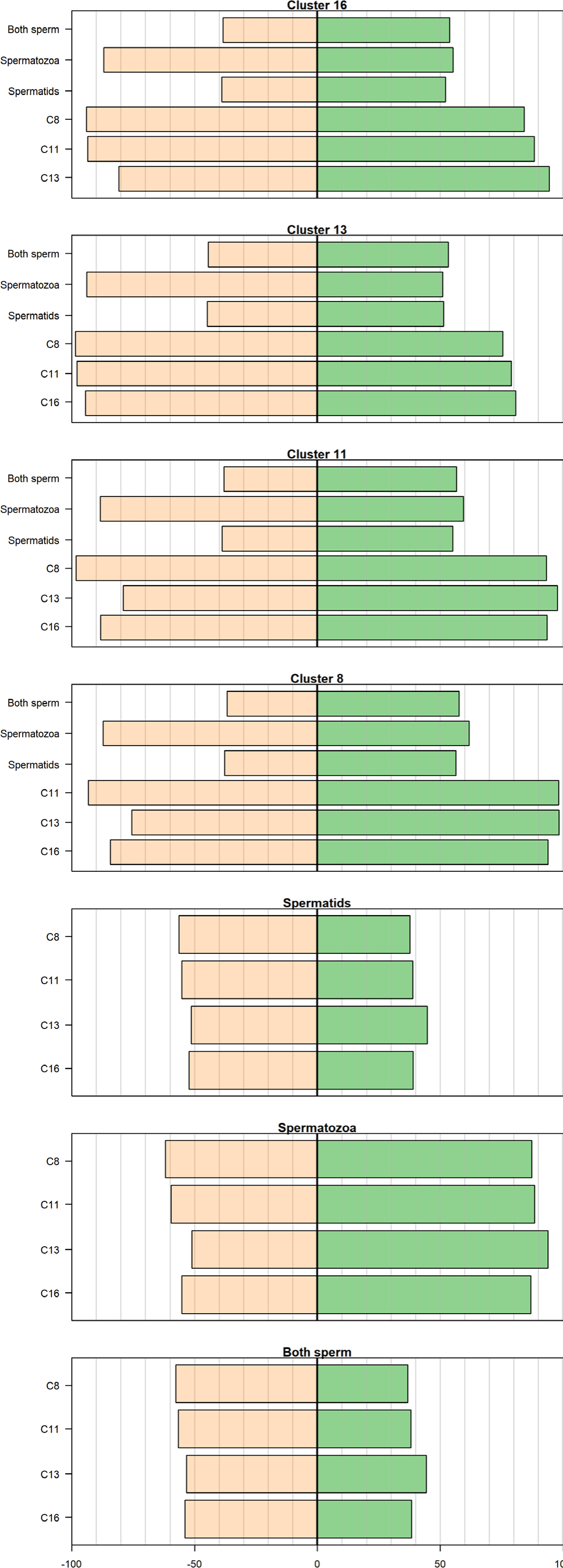
Proportions of features shared among putative sperm and oocyte/embryo clusters. In each panel green bars denote the percentage of features in the cluster labelled on the left that are found in the cluster in the title, while orange bars show the percentage of features in the cluster in the title that are found in the cluster labelled on the left. Features were considered found in a cluster if their average cluster expression value was > 0.05, which corresponds to the top 52% of feature expression values.

**Figure S6.**
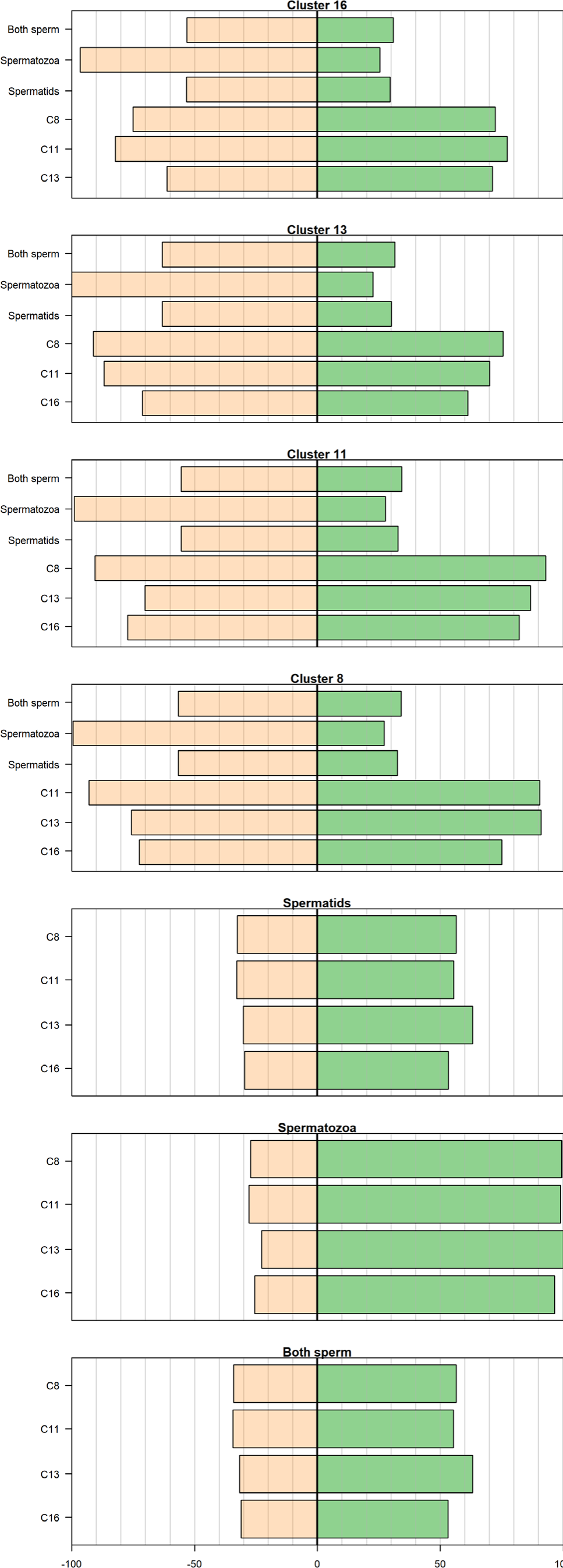
Proportions of features shared among putative sperm and oocyte/embryo clusters. In each panel green bars denote the percentage of features in the cluster labelled on the left that are found in the cluster in the title, while orange bars show the percentage of features in the cluster in the title that are found in the cluster labelled on the left. Features were considered found in a cluster if their average cluster expression value was > 0.6, which corresponds to the top 10% of feature expression values.

